# A mechanistic model of protein kinase A dynamics under pro- and anti-nociceptive inputs

**DOI:** 10.64898/2026.02.12.705506

**Authors:** Polina Lakrisenko, Jörg Isensee, Tim Hucho, Daniel Weindl, Jan Hasenauer

**Author notes:** These authors contributed equally (Polina Lakrisenko, Jörg Isensee). These authors contributed equally (Tim Hucho, Daniel Weindl, Jan Hasenauer).

## Abstract

Protein kinase A (PKA) is a central integrator of nociceptive signaling, yet a quantitative account of how pro- and anti-nociceptive inputs shape its dynamics remains incomplete. Here, we develop a mechanistic model of PKA activity in nociceptive neurons that explicitly links receptor activation to downstream kinase regulation. Using time-course and dose-response measurements, we infer unknown process parameters and quantify parameter and prediction uncertainties to ensure robust conclusions. The model captures the activation of PKA by serotonin and forskolin and its suppression by opioids. We show how the model can be used for the assessment of alternative circuit topologies, and demonstrate that receptor context and stimulation history reconfigure PKA responsiveness, providing testable predictions for opioid modulation under clinically relevant dosing. This framework offers a principled basis for integrating PKA with broader pain-signaling networks, supports rational exploration of combination therapies, and establishes a general strategy for disentangling neuromodulatory control of kinase activity.

**Author summary:** Pain perception is modulated by a complex network of signaling pathways activated by different receptors with opposing effects. A key player in this process is protein kinase A (PKA), whose regulation by both serotonin and opioid receptors is not yet fully understood. In this study, we developed a mathematical model to investigate how these opposing signals affect PKA activity in sensory neurons. After estimating the unknown model parameters from a comprehensive dataset, we were able to quantitatively analyze the dynamic behavior of the system and use it for comparison of alternative circuit topologies. Our model provides a valuable tool for integrating diverse molecular interactions involved in pain processing and could help guide future efforts to develop better treatments for chronic pain and reduce opioid tolerance.

## Introduction

Pain is a common feature of many disorders and, while initially protective, it can lose its value and become a pathological condition in itself [1, 2]. In some cases, pain progresses to a chronic state, persisting beyond the resolution of the underlying cause. Chronic pain not only exacerbates patient suffering, but also imposes a substantial burden on healthcare systems and society at large [3–5].

Opioids are widely used in the treatment of pain, particularly in acute and cancer-related cases [6–8]. Despite their long history of clinical use and extensive research, significant challenges remain—most notably the development of tolerance and the risk of dependence [4, 9]. One factor that hinders progress in addressing these challenges is that opioid-induced signaling is still not fully understood. This is due to complex interactions with other signaling pathways, resulting in a context-specific response. Among these interactions, the interplay between serotonin-mediated (serotonergic) and opioid-mediated (opioidergic) mechanisms represents a key component of pain signaling and a major source of complexity in neural dynamics [9–11]. The corresponding pathways exhibit context-dependent, tissue-specific effects, which can be either synergistic or antagonistic [9–11]. In the peripheral nervous system, for example, serotonergic receptors exhibit pro-nociceptive (pain-promoting) activity, counteracting the anti-nociceptive effects mediated by opioid receptors [12].

Pain reception takes place in specialized sensory neurons called nociceptors, whose cell bodies are located in the dorsal root ganglion (DRG) [1]. They are responsible for detecting potentially harmful stimuli and initiating the sensation of pain. This process is regulated by opposing action of different receptor systems, most notably by members of the G-protein-coupled receptor (GPCR) family [1, 13]. The G proteins are heterotrimers composed of *α, β*, and *γ* subunits. They are classified into distinct families based on their signaling effects and downstream targets, including stimulatory G proteins (G_S_) and inhibitory G proteins (G_I_).

Serotonergic and opioidergic mechanisms in the peripheral nervous system are represented by different types of receptors, including serotonin (5-hydroxytryptamine, 5-HT) receptors and *µ*-opioid receptors (MOR), both of which are members of the GPCR family. Specifically, 5-HT_4_ receptors expressed on nociceptors are coupled to G_S_ [10, 14], while MORs are coupled to G_I_. These receptors are activated by ligand binding and may undergo desensitization with prolonged stimulation, followed by potential resensitization through recycling or dephosphorylation [15, 16]. Receptor subtypes can differ in their constitutive activity, desensitization and internalization sensitivity, and ligand-binding affinities [11, 15, 17]. 5-HT_4_ and MOR receptors exert opposing effects on adenylate cyclase (AC). AC is an integral plasma membrane enzyme with its active site facing the cytosol, where it catalyzes the conversion of adenosine triphosphate (ATP) to cyclic adenosine monophosphate (cAMP), a second messenger responsible for propagating extracellular signals to intracellular targets [18].

Upon receptor activation, a conformational change promotes the exchange of GDP for GTP on the G*α* subunit, leading to its dissociation from the G*βγ* complex. The G*α* subunit, depending on its type, modulates downstream signaling: G*α*_S_ stimulates, while G*α*_I_ inhibits, cAMP production. cAMP activates protein kinase A (PKA), a holoenzyme composed of two regulatory (R) and two catalytic (C) subunits [18]. In its inactive state, the regulatory subunits bind and inhibit the catalytic subunits. Upon binding of four cAMP molecules to the regulatory subunits, the catalytic subunits are released and become active [19]. These free catalytic subunits then phosphorylate target proteins, such as ion channels, thereby modulating pain signaling pathways [1]. Additionally, the catalytic subunits can translocate to the nucleus, where they phosphorylate the cAMP response element-binding protein (CREB). Phosphorylated CREB regulates gene transcription, further influencing long-term changes in pain processing [18]. In particular, PKA has been shown to contribute to pain sensitization, characterized by increased sensitivity to noxious (harmful) stimuli and, in some cases, the perception of pain in response to normally harmless stimuli [13, 20–22].

PKA has been extensively studied over the past decades, and substantial progress has been made in elucidating its activation mechanisms. The advances include rigorous structural analyses of the PKA holoenzyme that elucidate cAMP-driven allosteric conformational changes underlying activation [23, 24], the discovery that cAMP-activated R subunits modulate G*α*_I_-coupled receptor signaling [25], the identification of PKA-dependent modulation of HCN2 channels as a mechanism underlying inflammatory pain sensitization in nociceptive sensory neurons [21], a refined understanding of the sequence of PKA subunit phosphorylation and its activation by cAMP [19], and the demonstration that depolarization-induced PKA activation is mediated by Ca_V_1.2 channels [26]. However, important knowledge gaps remain, particularly in the context of pain signaling. Specifically, there is a lack of integrative analysis that combines known molecular interactions to assess their relative contributions to specific phenotypes. Moreover, the role of potential feedback loops is unclear, and a fully quantitative, systems-level understanding of PKA regulation is still missing.

Mechanistic mathematical modeling offers a powerful tool to formalize biological assumptions and combine current knowledge about a biological process. It is especially useful for the analysis of complex networks, where keeping track of species and their interactions becomes difficult. Mechanistic models can be used to test the consistency of available knowledge, for analysis of latent quantities, and to predict dynamic system behavior under novel conditions. Moreover, mechanistic models allow for the integration of heterogeneous datasets and their joint analysis, offering a comprehensive approach to understanding dynamic processes.

Mechanistic models are most commonly formulated as systems of ordinary differential equations (ODEs), which describe the dynamics of the underlying biochemical reaction networks. This modeling approach has been widely and successfully applied to a variety of biological processes, including the cell cycle [27], gene regulatory circuits [28–30], and intracellular signaling pathways [31, 32]. Mechanistic models have also been applied for the analysis of pain signaling [33]. Examples include models of neuron firing in the spinal dorsal horn [34–36] and an elaborate dynamic computational model of the human spinal cord [37]. A number of mechanistic models have been developed to investigate pain signaling at the cellular level. These include a model of DRG neurons with focus on the P2 family of surface receptors [38], a model of inflammation-induced sensitization in a mechanosensitive muscle nociceptor [39], a mechanistic mathematical model of PKA-II activity in primary sensory neurons [19], an ODE-constrained mixture model of NGF-induced Erk1/2 phosphorylation in primary sensory neurons [40], and a more flexible model of the same pathway that accounts for multiple levels of heterogeneity simultaneously [41]. However, due to the diverse tissue types and the complex processes involved in pain signaling, effective modeling requires case-specific approaches that focus on the characteristic interactions relevant to each context.

In this study, we developed a comprehensive ODE-based model of PKA activation and cAMP production in DRG that captures the key steps from serotonin receptor stimulation to the dissociation of the catalytic subunit, and integrates the opposing effects of pro-nociceptive 5-HT_4_ and anti-nociceptive MOR signaling. We calibrated the model on a comprehensive dataset and used it to analyze the possibility of a feedback between PKA and G_I_ protein, and to illustrate the effect of receptor internalization on cAMP levels, as a proxy for pain signal intensity.

## Results

### Mechanistic model of serotonin and opioid receptor signaling in dorsal root ganglia

To quantitatively analyze PKA activity in rat dissociated cultured DRG sensory neurons in response to pro- and anti-nociceptive signaling, we developed an ODE-based model of 5-HT-induced activation and MOR-induced inhibition of PKA activity (Fig. 1). To develop the model, we conducted a comprehensive literature survey to identify and extract relevant molecular interactions. The model captures ligand-dependent receptor activation, subsequent G-protein signaling, and its regulatory effects on AC activity and cAMP synthesis, ultimately linking these processes to PKA activation through cAMP binding. The description of the PKA activation through cAMP binding is based on our previous model [19], which we extended here to include GPCR and G-protein signaling. There are different isoforms of both R (RI*α*, RI*β*, RII*α*, and RII*β*) and C (C*α* and C*β*) subunits. Here, we focus on PKA composed of RII and C*α* subunits, as these isoforms are more relevant for the interactions that we consider [19]. Specifically, both RII isoforms localize to the cell membrane and play active roles in GPCR signaling. RII*β* in particular is expressed at high levels in nociceptors [42].

**Fig 1.**
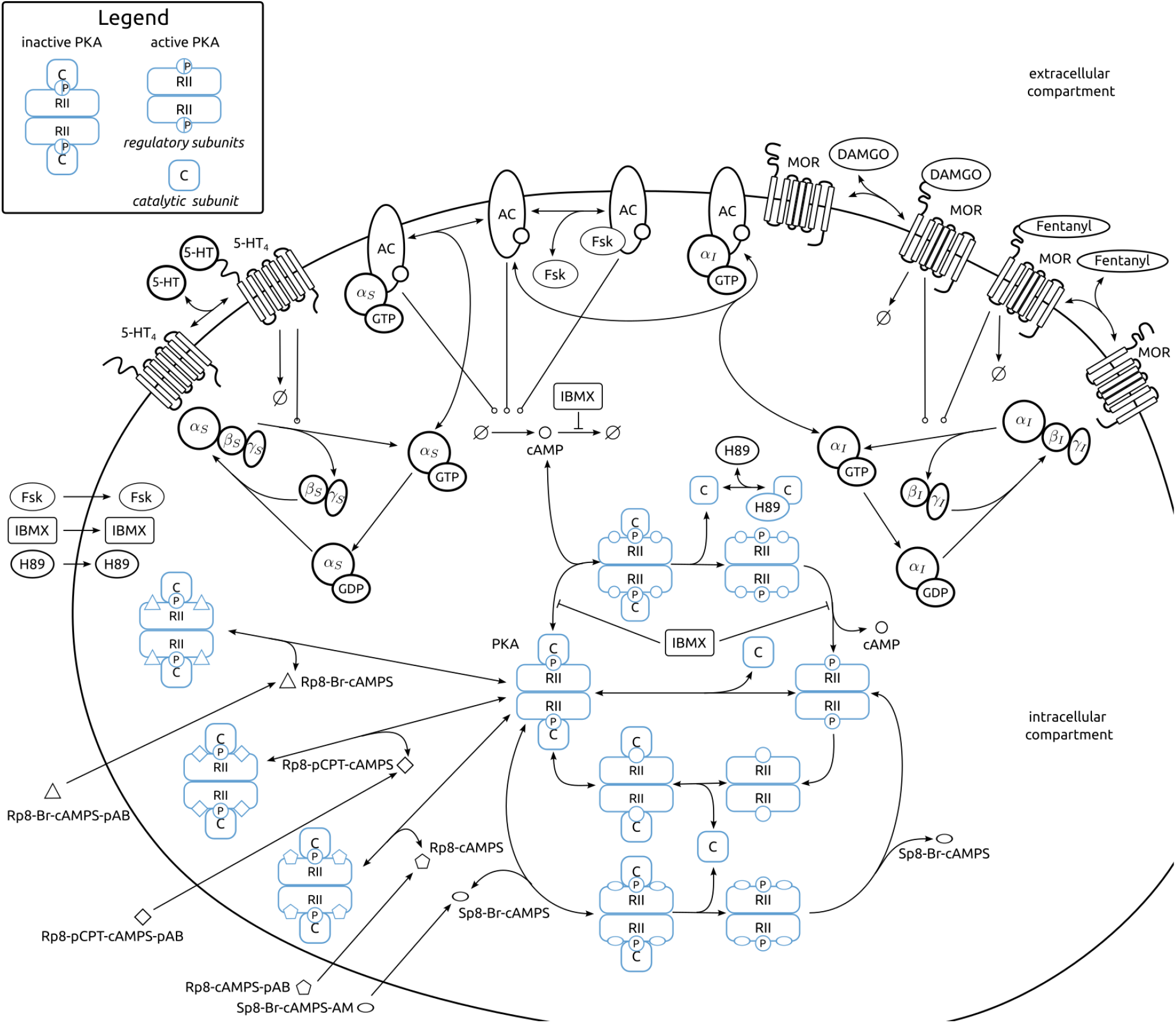
Model of PKA activity modulation by serotonin and opioid receptor signaling. In DRG, cAMP production is regulated by GPCRs such as 5-HT_4_ and MOR, which modulate AC activity through distinct G-protein signaling pathways. Upon ligand-induced receptor activation, GDP is exchanged for GTP on the G protein *α* subunit, followed by its dissociation from the G*βγ* complex. cAMP production is stimulated by *α*_S_ and inhibited by *α*_I_, which modulates PKA activity. When cAMP binds to the regulatory subunits (RII) of PKA, it induces a conformational change that releases the two active catalytic subunits (C). The degradation reaction of the ligand-bound receptors represents both receptor degradation and internalization. Blue borders indicate species accessible by the employed measurements techniques.

The developed model provides a description for a broad spectrum of perturbations with eight compounds: (1) forskolin (Fsk), which increases cAMP levels by stimulation of AC; (2) 3-isobutyl-1-methylxanthine (IBMX), which inhibits cAMP degradation; (3) H-89, which inhibits the C subunit, blocking its downstream activity and preventing its reassociation with the regulatory subunit; (4-7) two types of cAMP analogs, Sp- and Rp-isomers, which serve as activators and inhibitors of PKA, respectively; (8) 5-HT, which activates 5-HT_4_ receptors, leading to upregulation of cAMP production; and

(9, 10) opioids DAMGO and fentanyl, which activate MORs, leading to downregulation of cAMP production (Table 1). As core readouts, we consider the phosphorylated PKA regulatory subunit (pRII) and the catalytic subunit as indicators of PKA activity, given their central roles in downstream signaling processes.

**Table 1.** Overview of the different stimuli included in the dataset for model calibration and their mode of action.

| # | Compound | Action | Reference |
| --- | --- | --- | --- |
| (1) | Fsk | Activator of cAMP production | [12] |
| (2) | IBMX | Inhibitor of cAMP degradation | [19] |
| (3) | H-89 | Inhibitor of catalytic subunit | [19] |
| (4) | Sp-8-Br-cAMPS-AM | cAMP analog, PKA activator | [19] |
| (5) | Rp-cAMPS-pAB | cAMP analog, PKA inhibitor | [19] |
| (6) | Rp-8-pCPT-cAMPS-pAB | cAMP analog, PKA inhibitor | [19] |
| (7) | Rp-8-Br-cAMPS-pAB | cAMP analog, PKA inhibitor | [19] |
| (8) | 5-HT | G<sub>S</sub>-GPCR agonist | [12, 19] |
| (9) | DAMGO | G<sub>I</sub>-GPCR agonist | [12] |
| (10) | Fentanyl | G<sub>I</sub>-GPCR agonist | [12] |

Overall, the model includes 38 interacting species and 58 reactions in the cytosol, and depends on 76 unknown parameters. We parameterized the rates of (reversible) complex formations in terms of the rate constant of the forward reaction (k_f_) and the dissociation constant (K_D_) values. For related reactions that may exhibit slight differences in kinetics, such as those arising from different phosphorylation states of the substrates or from distinct activation states due to ligand binding or absence, we introduced multiplicative modifiers (*ξ*) to adjust the relevant parameters for each specific reaction. The import rates and receptor internalization rates were parameterized using proportionality factors (k_i_ and k_int_, respectively). The unknown parameters consist of 55 model parameters, such as reaction rate and dissociation constants, and 21 observation model and noise parameters. The model is implemented in the Systems Biology Markup Language (SBML) [43], facilitating reuse and extension.

### Calibrated model accurately reproduces all experimental conditions

To evaluate the validity of the proposed model, we assessed its ability to describe experimental data. Therefore, we assembled a comprehensive and coherent dataset from rat DRG sensory neurons from results published in [12] and [19]. These publications provide high-content screening (HCS) immunofluorescence microscopy measurements of pRII and C_*α*_, and immunoblotting measurements of activation-induced changes in RII phosphorylation in cell lysates (S1 Text, Fig. S1). In total, the dataset comprises 14 experimental panels, with overall 211 conditions (controls, individual compounds and their combinations). Altogether, it contains 2,121 measurements of phosphorylated RII (pRII) and 42 measurements of C_*α*_ in populations of primary sensory neurons (S1 Text, Table. S1). This dataset has been split in training and validation data following the suggestion of the experimental partners. In addition, we collected prior information on several parameters, including dissociation constants for AC and Fsk, as well as proportionality constants describing the differences in PKA activation and deactivation rates when RII is phosphorylated versus unphosphorylated (S1 Text, “Parameter priors” Section). The data and metadata are encoded in the PEtab format [44] and shared via the Zenodo repository (https://doi.org/10.5281/zenodo.18619229) to facilitate their use in future studies.

To determine the unknown model parameters, we performed maximum a posteriori (MAP) estimation using multi-start local optimization [45]. As the estimation problem is high-dimensional and computationally challenging, we employed an informed start point sampling algorithm (see S1 Text, “Parameter starting point sampling” Section). We sampled the starting points for the 46 parameters in the PKA activation model from log-normal distributions which possess as median the estimates reported by [19]. The remaining 30 parameters were initialized using log-uniform sampling within predefined bounds. In total, we performed 7,000 local optimizations.

The analysis of the optimization results indicates a good reproducibility of the fitting results, with the differences among the 37 best objective function values not being statistically significant (Fig. 2a), as evaluated using a *χ*^2^ distribution (see Materials and Methods section). Overall, the simulated observables are in good agreement with the training data for the MAP estimate (Fig. 2b). This agreement is further supported by comparisons between model simulations and experimental data from [12] and [19]. The model reproduces key qualitative features of the measurements, including the dose-dependent inhibition of pRII induction by Rp-analogs of cAMP (dataset 2), the dose-dependent opposing effects of forskolin and H-89 on signal intensity (dataset 4), and the attenuation of the signal by opioids (datasets 7, 8, 10, and 11).

**Fig 2.**
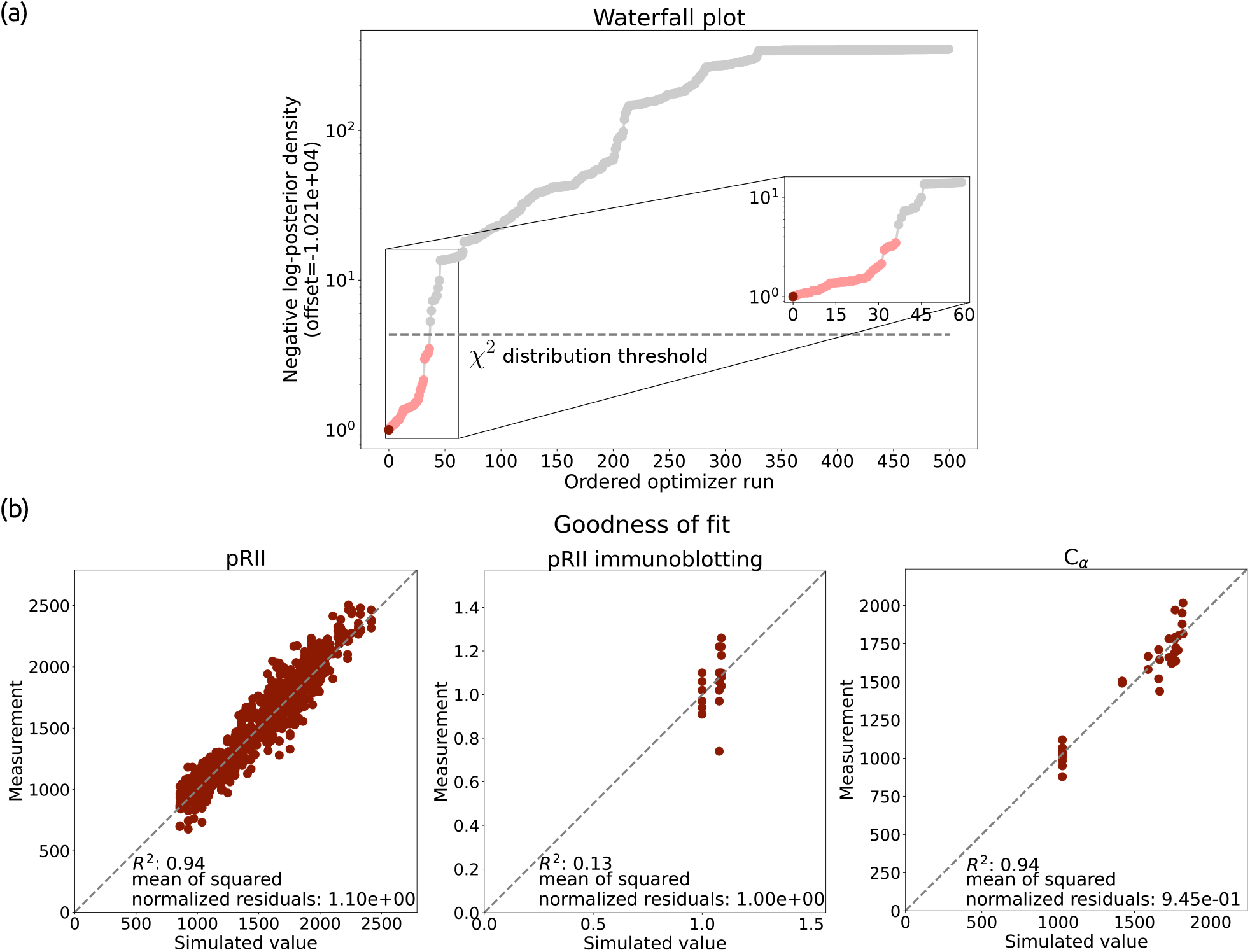
Optimization results. (a) Waterfall plot of the best 500 of 7,000 local optimizations, and zoom in of the best 60. The dashed line indicates a threshold based on the 99th percentile of a *χ*^2^ distribution with 1 degree of freedom. The optimizations below the threshold are indicated by the red color shades. The dark red indicates the MAP estimate. (b) Scatter plot for the experimental data and model simulations for the MAP estimate.

**Fig 3.**
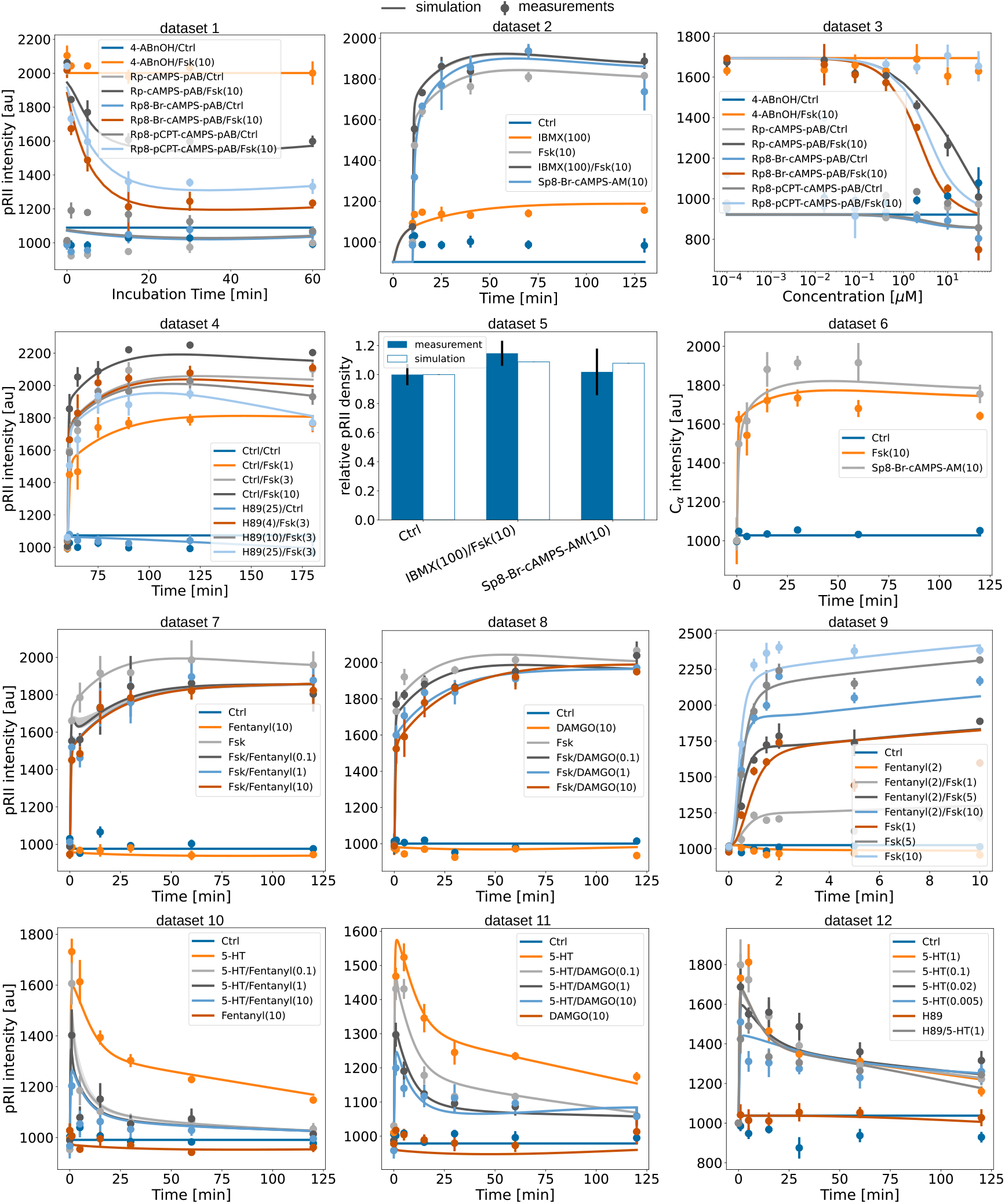
Optimization results. Line plots and a bar plot for measurements and model simulations using an ensemble of parameter sets from the best optimization runs, selected based on the objective function value threshold corresponding to the 99th percentile of a *χ*^2^ distribution with 1 degree of freedom. The error bars indicate the standard deviation computed from measurement replicates.

Notably, the simulations capture the more pronounced dose response of DAMGO (datasets 8 and 11) compared with fentanyl (datasets 7 and 10). Although some experiments are not perfectly reproduced (e.g., dataset 12), these discrepancies can be partially attributed to the high level of noise in the experimental data.

Overall, the model provides a very accurate description of the data for all experimental panels, capturing in total time-resolved data for 162 conditions (control condition, single compound treatments and combination treatments). This is the case despite the high-dimensional parameter space and the apparently complex objective function landscape, reflected in a relatively low convergence rate of the multi-start optimization method.

### Parameter and prediction uncertainty

To assess the parameter estimate and the model predictions, we performed a comprehensive uncertainty analysis. We determined the parameter uncertainties by computing profile posteriors and deriving credible intervals [46] (see Materials and Methods). The analysis revealed that 34 out of 55 model parameters (61.8%) are practically identifiable, i.e. have finite 99% credible intervals (within the parameter bounds) (Fig. 4). The inspection of the positioning of identifiable and non-identifiable parameters in the network revealed that all parameters related to the dynamics of PKA are identifiable. In contrast, 71% of parameters related to G protein dynamics are practically non-identifiable (Fig. 4). This is not unexpected as the G protein level dynamics is not directly observed. Moreover, for reversible, joint estimation of the forward rate and the dissociation constant is often challenging, as also observed here.

**Fig 4.**
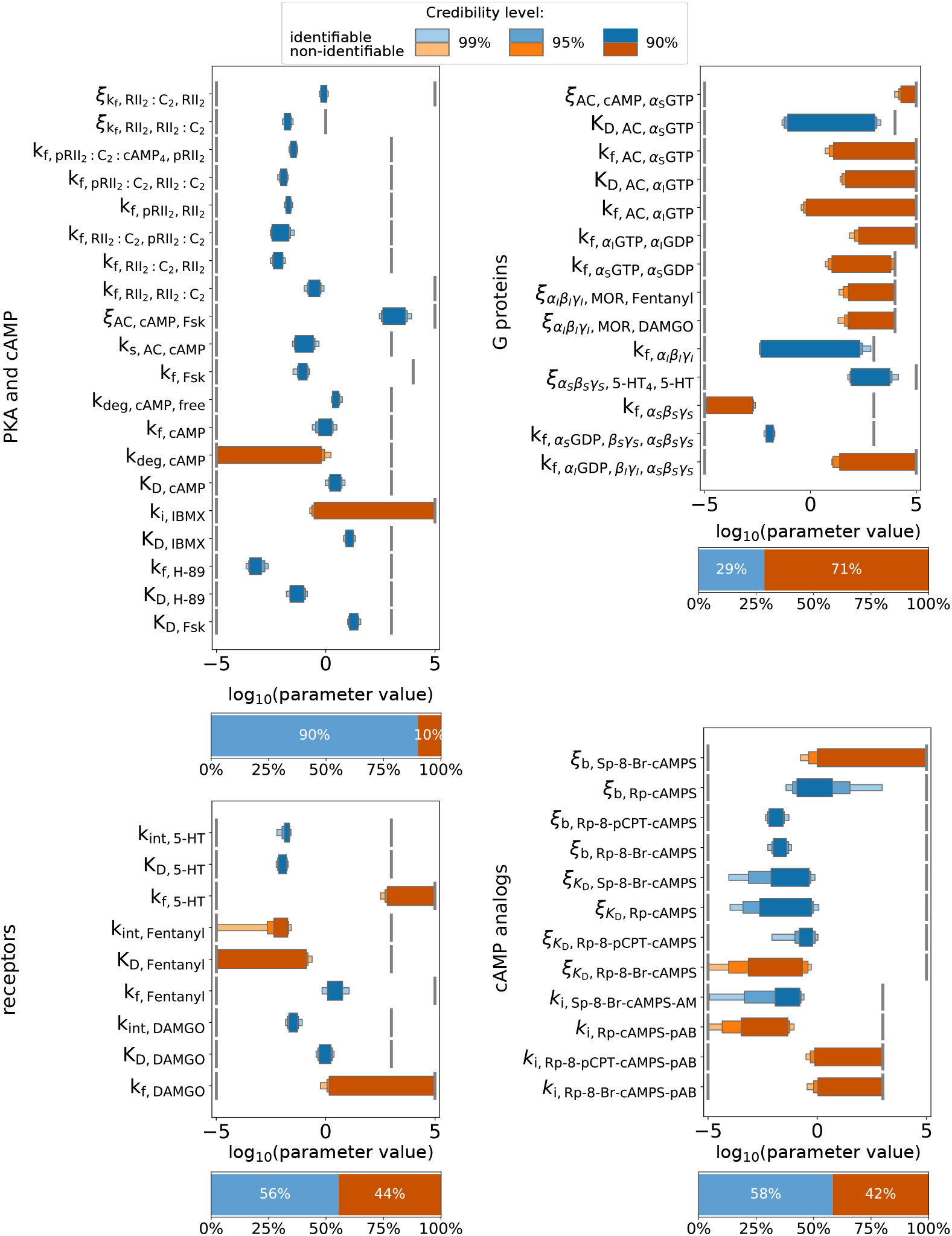
Parameter uncertainty. Credible intervals of the model parameters computed using profile posteriors. The intervals for identifiable parameters are shown in blue shades, for non-identifiable parameters in orange shades. Vertical gray lines indicate the parameter bounds used for optimization. Stacked bar plots show the fraction of identifiable and non-identifiable parameters within each group.

For most binding and dissociation constants, namely for 5-HT, Fentanyl, DAMGO, *α*_*S*_GTP and *α*_*I*_ GTP, at least one of the two is non-identifiable. Both parameters are identifiable only for the binding of cAMP to PKA, H-89 to the regulatory subunits and Fsk to AC (Fig. 4).

In order to assess the prediction uncertainties, we used an ensemble-based approach [47]. Therefore, an ensemble of parameter vectors was assembled from the parameter vectors obtained during parameter optimization. The ensemble contains parameter vectors that achieve fitting results statistically indistinguishable from the MAP estimate at a significance level of 0.01, corresponding to a difference of 3.32 in the objective function value (see Materials and Methods). The resulting parameter ensemble contains 2,877 parameter vectors and is substantially more diverse than the set of 37 optimization results considered above (S1 Text, Fig. S3).

The ensemble-based analysis of the prediction uncertainties reveals that the uncertainties in the observables are minimal (Fig. 5). Yet, prediction uncertainties for several state variables are large, particularly those involving AC and its associated complexes. This uncertainty pattern is consistent with the pattern of the parameter non-identifiabilities. Interestingly, the shape of the simulated state trajectories is consistent across most simulations in the ensemble, merely the scale differs (S1 Text, Fig. S3). Such scale uncertainty is not unexpected given the lack of quantitative measurements for these species and the fact that the measurements reflect aggregates of multiple state variables rather than individual ones (S1 Text, “Observation model Section”).

**Fig 5.**
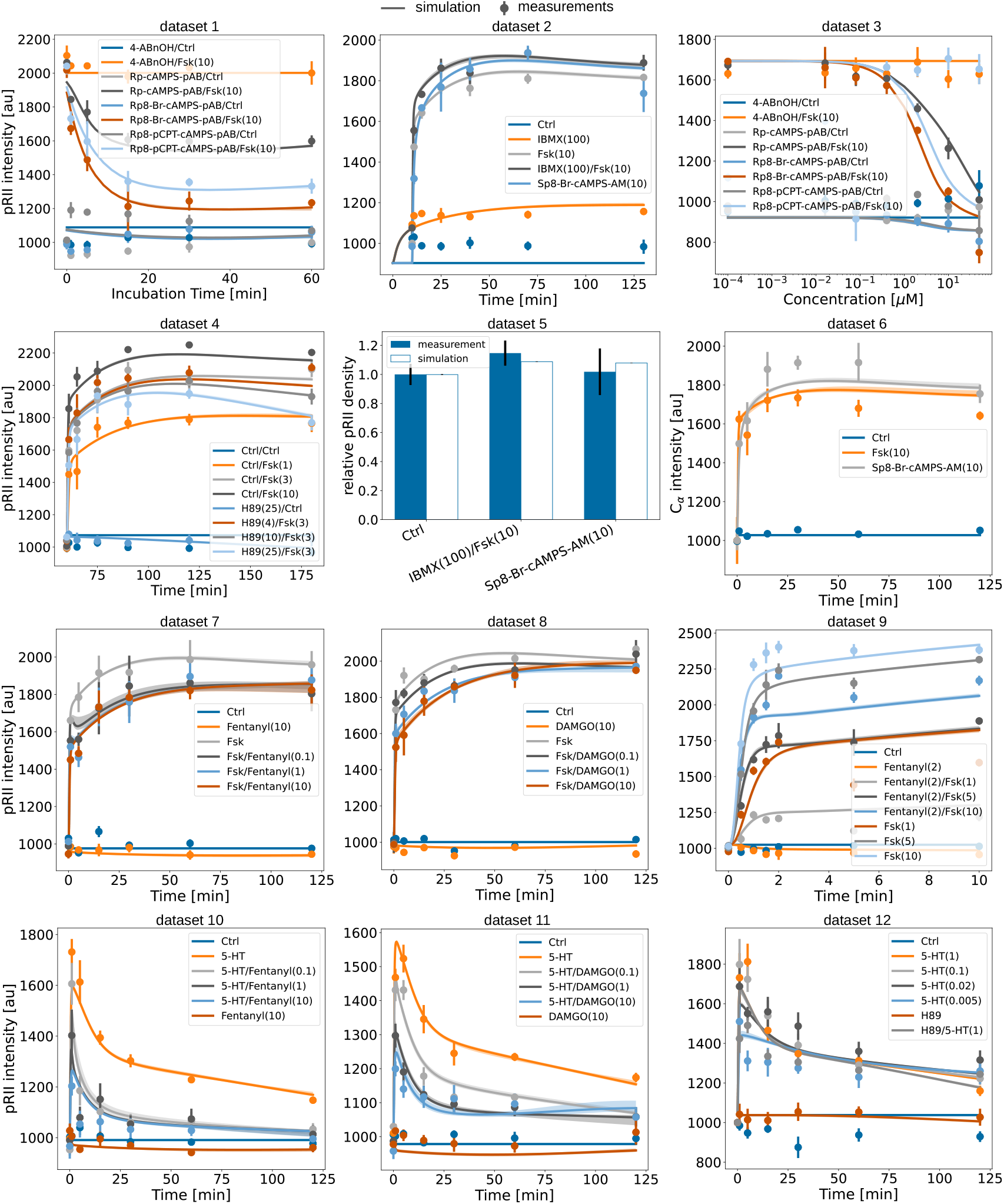
Prediction uncertainty. Prediction uncertainty was estimated using an ensemble of parameter sets from the best optimization runs, selected based on the objective function value threshold corresponding to the 99th percentile of a *χ*^2^ distribution with 1 degree of freedom, 2,877 parameter vectors in total. The error bars indicate standard deviation computed from measurement replicates.

Overall, the uncertainty analysis reveals that various parameters, state and observable characteristics are well determined by the data. Yet, the lack of more detailed quantitative measurements for individual species results in uncertainty of the scales.

### Model reliably predicts validation datasets

To validate the predictive power of the model, we used it to simulate experimental conditions that were not used for parameter estimation. The validation datasets comprise three independent experiments: (1) a comparison of pRII levels in response to 5-HT stimulation with and without IBMX, which inhibits cAMP degradation and thereby amplifies the signaling response; (2) measurement of pRII intensity following stimulation with varying doses of DAMGO or fentanyl; (3) pRII responses to stimulation with 5-HT, forskolin, or control, after different durations of pretreatment with either control or fentanyl.

We simulated the experimental conditions using the 2,877 parameter sets from the best optimization runs, selected based on the objective function value threshold corresponding to the 99th percentile of a *χ*^2^ distribution with 1 degree of freedom. The simulation demonstrated that the model can qualitatively predict the observed outcomes despite the presence of non-identifiable parameters and parameter uncertainties spanning several orders of magnitude (Fig. 6). The simulations reproduced the signal-enhancing effect of IBMX, but slightly overestimate its strength (Fig. 6a). The model also reproduced the overall opioid dose–response trend in the dataset 13; however, it did not fully reproduce the differences between DAMGO and fentanyl (Fig. 6b). Additionally, the simulations reflected the signal-dampening effect of fentanyl pretreatment on subsequent stimulation with both 5-HT and forskolin (Fig. 6c). The relative error between measurement and simulation is on average around 4%.

**Fig 6.**
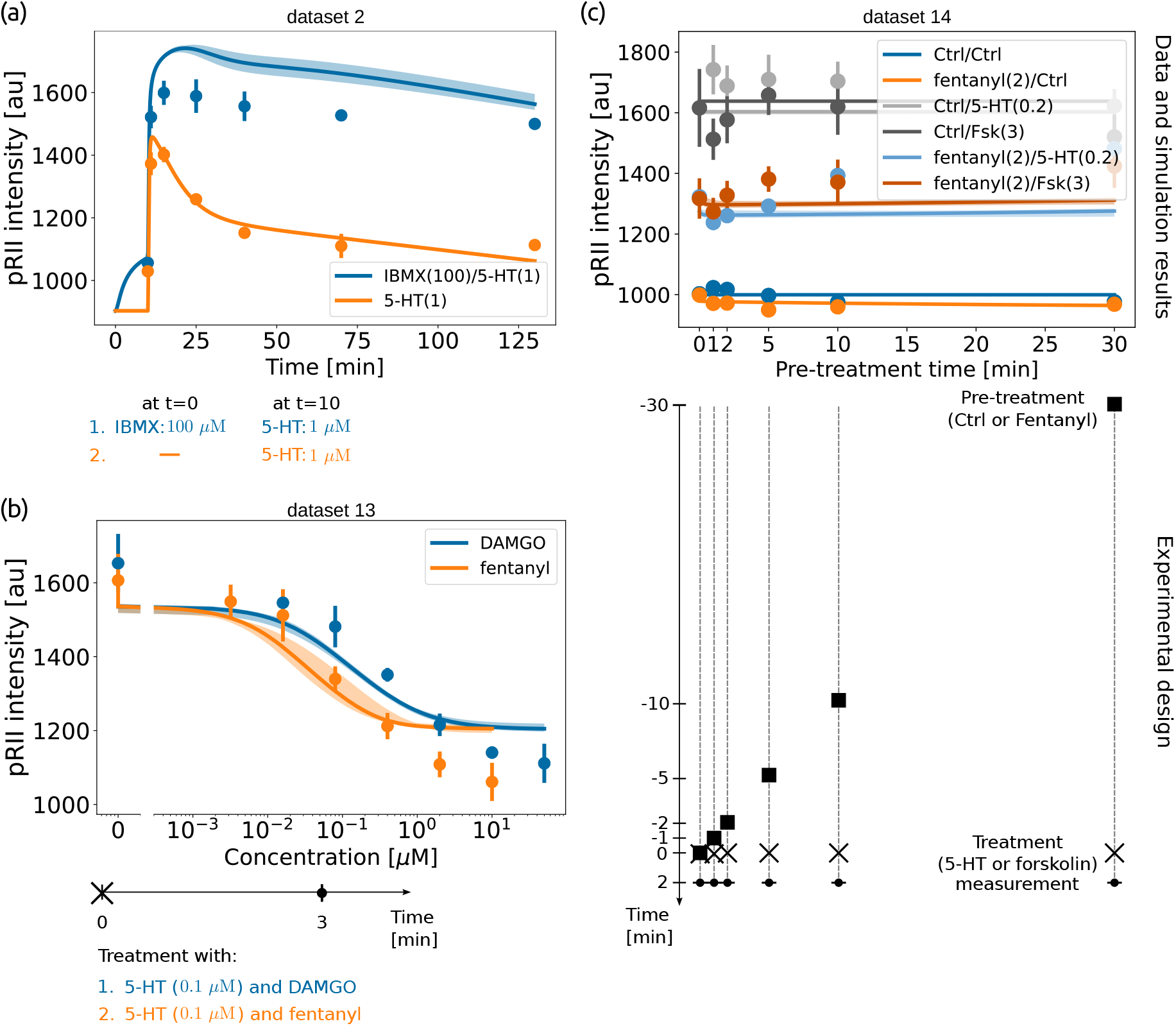
Model predictions compared to experimental results. Ensemble predictions using 2877 parameter sets from the best optimization runs, selected based on the objective function value threshold corresponding to the 99th percentile of a *χ*^2^ distribution with 1 degree of freedom. For each experiment, model predictions are compared with the data (top), and the experimental design is shown (bottom). (a) pRII response induced by 1 *µ*M 5-HT after 10 minute pretreatment with 100 *µ*M IBMX (blue) and induction of pRII intensity by 1 *µ*M 5-HT (orange). (b) Dose-response curves showing the effect of 0 to 50 *µ*M of DAMGO (blue) and of 0 to 10 *µ*M of fentanyl (orange) after stimulation with 0.1 *µ*M 5-HT. (c) pRII intensity in response to stimulation with 0.2 *µ*M 5-HT, 3 *µ*M of forskolin, or control after 0, 1, 2, 5, 10 or 30-minute pretreatment with control or 2 *µ*M of fentanyl. The error bars indicate standard deviation computed from measurement replicates. The shaded area represents the range of all ensemble simulations.

Overall, the results demonstrate that the model can be used for meaningful *in silico* experiments and underscore its ability to provide reasonable prediction.

### Model illustrates receptor desensitization due to internalization

The proposed model provides a mechanistic description for a broad spectrum of datasets for different stimuli. Here, we used this to study the interplay between pro- and anti-nociceptive signaling as well as receptor internalization dynamics. To account for the parameter uncertainties, we used the full parameter ensemble.

To assess the difference between MOR activation by DAMGO and fentanyl, alongside activation of 5-HT receptors, we performed simulations with varying concentrations of 5-HT (Fig. 7a). The ensemble simulations indicate concentration-dependent differences in the response amplitude: the higher the 5-HT concentration, the higher the predicted maximal pRII concentration, which illustrates pro-nociceptive activity of 5-HT receptors (Fig. 7a). Although simulations show greater uncertainty with fentanyl co-treatment than with DAMGO (Fig. 7b), fentanyl’s higher potency is clearly evident in the lower values following the initial peak and in the persistently lower values thereafter. Interestingly, at later time points, lower 5-HT concentrations resulted in higher sustained pRII levels. This behavior can be attributed to receptor internalization dynamics. Since in our model only ligand-bound receptors are internalized (Fig. 1), higher concentrations of 5-HT promote increased formation of 5-HT_4_:5-HT complexes, which are subsequently internalized, reducing the number of surface receptors available for continued signaling over time, independently of the co-treatment with either DAMGO or fentanyl (Fig. 7c). As a result, the simulations suggest that higher ligand concentrations elicit a rapid and strong initial response that is followed by a decline in activity due to receptor internalization, eventually leading to receptor desensitization. In contrast, lower ligand concentrations produce a less acute but more sustained signaling response.

**Fig 7.**
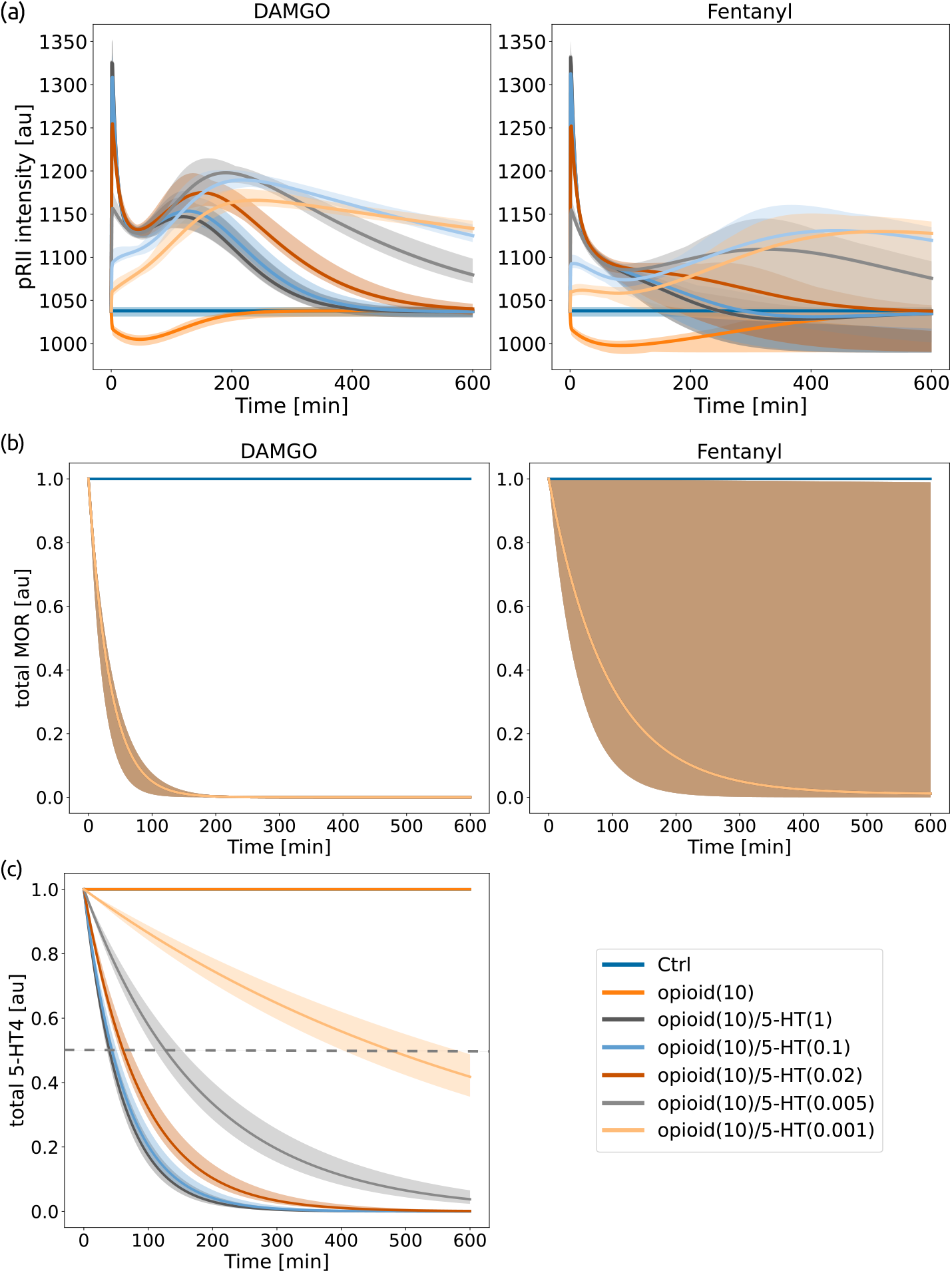
*In silico* experiment simulating MOR activation and concurrent activation of 5-HT receptors using varying concentrations of 5-HT. (a) Simulation of pRII microscopy measurements. Addition of 10*µ*M of DAMGO (left) or of fentanyl (right) at *t* = 0 was simulated. (b) Total amount of MOR over time, including both unbound and ligand-bound forms, relative to the amount at *t* = 0. (c) Total amount of 5-HT_4_ receptor over time, including both unbound and ligand-bound forms, relative to the amount at *t* = 0. This figure holds for both opioids as 5-HT_4_ dynamics is only influenced by 5-HT concentration. The dashed line indicates 50% of initial receptor amount. The shaded area represents the range of all ensemble simulations.

To assess how well the model predictions align with previous findings, we compared the rate of receptor internalization obtained from the model simulations with the estimates reported in [11]. According to the simulations, 50% of 5-HT_4_ receptors are internalized approximately 40 minutes after stimulation with 1 *µ*M 5-HT. This agrees well with previous observations, which reported pronounced internalization of the 5-HT_4*b*_ splice variant after 42–49 minutes of incubation with 1 *µ*M 5-HT [11].

Overall, the simulation study shows that the model can be used to assess properties of the process which are challenging to assess experimentally. Furthermore, it enables the assessment of emerging properties shaped by the interplay of different processes, such as peak height and duration.

### Model-based analysis of the comprehensive PKA data facilitates hypothesis testing

It has been reported that cAMP-bound R subunits enhance ligand-triggered 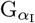 activation, leading to increased amplitude and duration of anti-nociceptive signaling [25]. This mechanism establishes a negative feedback loop between pro-nociceptive and anti-nociceptive signaling. To date, this feedback has been demonstrated primarily in kidney-derived cell lines, and its relevance in sensory neurons remains unclear. Our core model does not incorporate this feedback mechanism. To test whether the data by [12] and [19] indirectly supports such a mechanism, we extended the model by an additional reaction for G_I_ protein activation, with a rate proportional to the concentration of the pRII_2_:cAMP_4_ complex (Fig. 8a). This reaction depends on an additional reaction rate constant 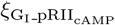, which had to be estimated.

**Fig 8.**
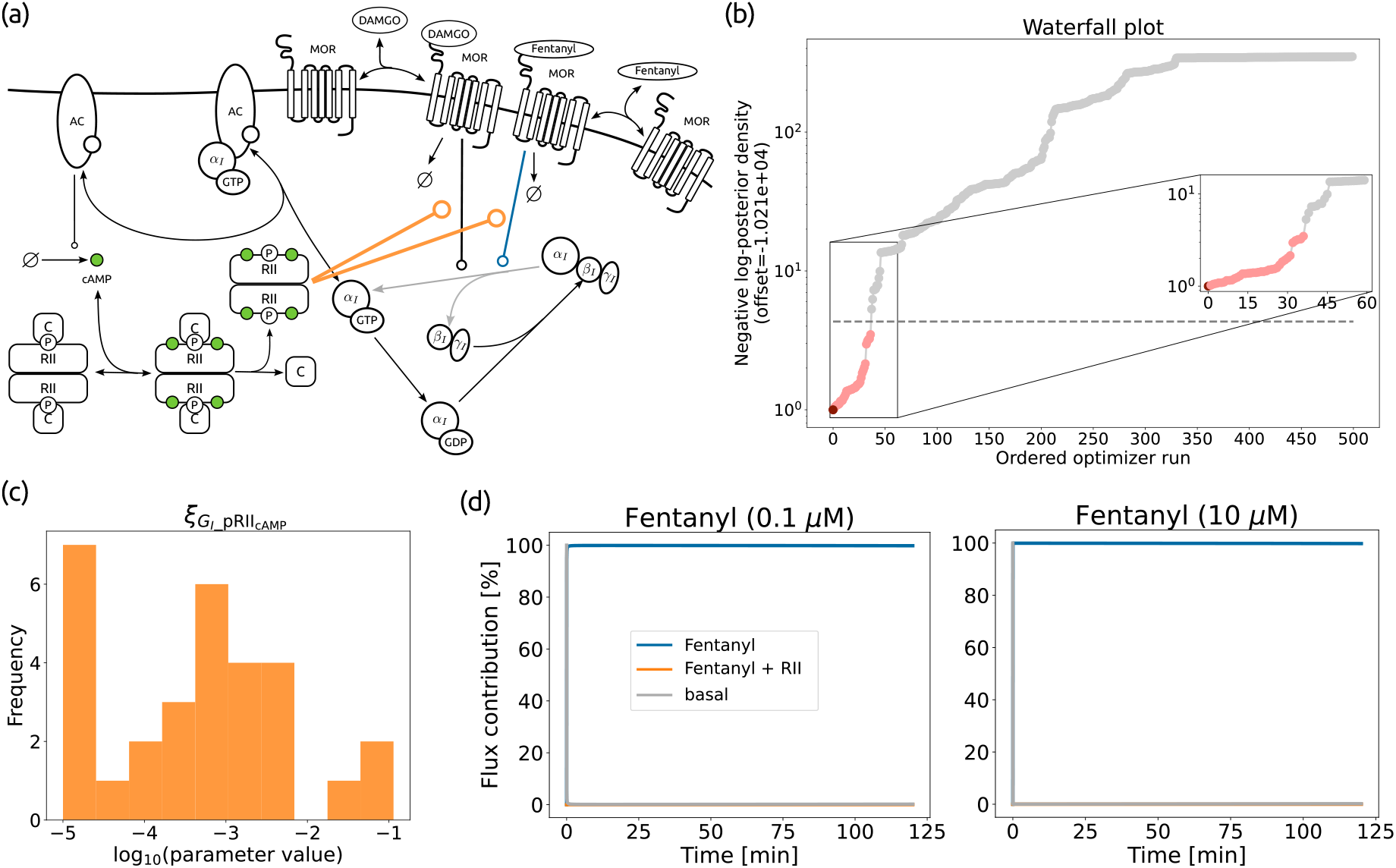
Assessment of potential cAMP-dependent interaction between G-protein and RII. (a) Alternative model structure incorporating the influence of cAMP-bound R subunits on 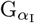 activation. In addition to G_I_ protein activation modulated by MORs stimulated either DAMGO or fentanyl (denoted by the lines ending a circle, black and blue, respectively) we add the suggested feedback mechanism (denoted by the orange lines ending in a circle). (b) The waterfall plot of the best 500 of 7,000 multi-starts, ordered by log-posterior value. The dashed line indicates a threshold based on the 99th percentile of a *χ*^2^ distribution with 1 degree of freedom and the red points the statistically indistinguishable optimization runs. (c) Histogram of the estimated values of the 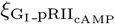 parameter for the 30 best optimization runs. (d) Flux contribution to the G_I_ protein activation over time after stimulation with 10 *µ*M of forskolin and different concentrations of fentanyl: basal (gray), modulated by fentanyl (blue), and by both fentanyl and the pRII_2_:cAMP_4_ complex (orange). Predicted using parameter vectors corresponding to the 30 best optimization runs below the *χ*^2^-based threshold.

We calibrated the modified model using the training dataset to investigate whether it improves the fitting results or leads to better model predictions. The reproducibility of the results was reasonable, yielding 30 parameter vectors that are statistically indistinguishable from the MAP estimate (as previously, assessed using the threshold based on the *χ*^2^ distribution, Fig. 8b). However, the model with the additional regulatory mechanism yielded a slightly worse objective function value than the original model. The resulting differences in information criteria (ΔAIC = 2.1; ΔBIC = 7.55) were within the commonly accepted threshold and are therefore not considered substantial (Table 2). This indicates that the two models are not statistically distinguishable and perform similarly in terms of overall fit on both training and validation datasets. Moreover, the estimated values of 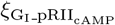 for the 30 best optimization runs are consistently small (Fig. 8c). Correspondingly, the relative flux contribution of the reaction representing the hypothesized mechanism is minimal (Fig. 8d). Hence, the current dataset does not provide sufficient evidence that the rate of G_I_ activation is dependent on the concentration of the pRII_2_:cAMP_4_ complex.

**Table 2.** Model comparison.

| Model | $n_\theta$ | $-\log \mathcal{L}_{\mathcal{D}}$ | AIC | BIC |
| --- | --- | --- | --- | --- |
| original (no feedback) | 76 | 10208.78 | 20569.56 | 20984.21 |
| model with the feedback | 77 | 10208.83 | 20571.66 ($\Delta = 2.1$) | 20991.76 ($\Delta = 7.55$) |

Overall, this study demonstrates that the model provides a useful platform for *in silico* hypothesis testing, complementing laboratory experiments. It enables systematic evaluation of the consistency of proposed regulatory mechanisms and helps determine whether available data are sufficient to support or refute specific mechanistic assumptions.

## Discussion

A comprehensive and quantitative understanding of pain signaling, including processes such as pain sensitization, requires integrative modeling approaches capable of capturing the complexity of diverse and sometimes opposing cellular mechanisms [33]. In this study, we developed a novel model of PKA activity in DRG sensory neurons modulated by serotonin receptors (5-HT_4_) and *µ*-opioid receptors, which respectively activate and inhibit PKA. To our knowledge, this represents one of the most detailed cellular-level models of pain signaling to date. We estimated 76 unknown model parameters using a comprehensive training dataset comprising 1,730 experimental measurements across 162 distinct conditions. The model accurately reproduces the available experimental data and enables *in silico* experiments, providing access to intracellular dynamics and quantities that are not directly measurable in experiments.

We used the calibrated model to perform an *in silico* experiment investigating the interplay between pro-nociceptive and anti-nociceptive signaling. The simulation highlights the impact of 5-HT_4_ receptor internalization on pain signaling intensity, showing that high ligand concentrations can trigger a rapid initial response, followed by a decline in activity due to reduced receptor availability. Yet, while our model provides valuable insights into 5-HT_4_ receptor signaling, many open questions remain. Overall, the dynamics of 5-HT_4_ receptors is less well studied compared to those of MORs. Studies have shown that these dynamics can vary significantly depending on the splice variant, and multiple variants may be expressed within the same cell type [10, 11, 17]. The 5-HT_4*b*_ splice variant interacts with G_I_ proteins in addition to G_S_ [11], a feature not captured by our current model. The 5-HT_4*a*_ isoform was shown to display little recycling compared to 5-HT_4*b*_ and to exhibit more prominent constitutive internalization [15, 17]. Currently, it is not clear which isoforms are more prevalent in DRG neurons. Receptor internalization may contribute to the development of drug tolerance and is crucial for determining appropriate dosing regimens. Therefore, further investigation of receptor dynamics is necessary for more detailed modeling and better understanding of 5-HT_4_ receptor activity.

Our model enabled the investigation of a potential negative feedback loop between PKA and G_I_ protein activity. However, model selection did not provide support for the presence of this feedback based on the available dataset. Importantly, these results do not rule out the possibility that such regulation may become relevant under alternative experimental conditions. Additional experiments, particularly those directly probing the interaction between the PKA regulatory subunit and the G_I_ proteins in sensory neurons, would therefore be valuable to further assess the physiological relevance of this mechanism.

As highlighted in previous work [48], a key challenge in understanding pain signaling lies in integrating diverse cellular processes, such as receptor activation, internalization, recycling, and downstream intracellular signaling, into cohesive frameworks. Our model addresses this need by providing a foundational component that can be extended to build more comprehensive models of pain signaling and drug response. For example, it can be further extended to explicitly include downstream effectors involved in pain sensitization, such as CREB, which regulates gene expression, and extracellular signal-regulated kinase (ERK), which modulates ion channel activity. More detailed description of both serotonin and MOR receptor internalization and recycling would enable investigation of potential feedback loops, such as G_*βγ*_ mediated MOR recycling [49] or its regulation by cAMP levels [50], and their impact on receptor desensitization. Moreover, it can be integrated with existing pharmacokinetic-pharmacodynamic models of opioids to support the development of optimized dosing strategies [48].

The model we present provides a valuable step toward integrative systems-level analysis of pain signaling and opioid pharmacology. By integrating known molecular interactions, it enables quantitative assessment of their relative contributions to PKA regulation, a central component of nociceptive signaling. Furthermore, the model lays the groundwork for exploring potential feedback loops and supports the development of a systems-level understanding needed to inform strategies for minimizing opioid tolerance and improving chronic pain management.

## Materials and Methods

### Mechanistic modeling

To describe the dynamic interactions of the species of interest (Fig. 1), we formulate an ordinary differential equation model of the form

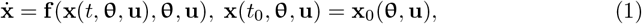

where the vector field 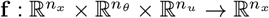, describing reaction stoichiometry and kinetics, determines the evolution of a vector of state variables 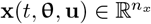, representing the concentrations of biochemical species, in time, denoted by *t* ∈ ℝ. The vector 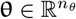 denotes unknown parameters, and the vector 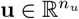 is known inputs that specify experimental conditions by directly specifying model component values that represent the external stimuli summarized in the Table 1. The vector field of the ODE model is assumed to be Lipschitz-continuous with respect to **x**, and the initial condition at *t* = *t*_0_ is 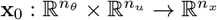.

We assume that the system (1) is in a steady state at the initial time point. In other words, the initial condition is defined as the steady state corresponding to a pre-equilibration input **u**^*e*^: **x**_0_(**θ, u**) = **x**^*^(**θ, u**^*e*^), where **x**^*^ is the steady state of

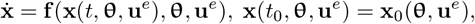

i.e.

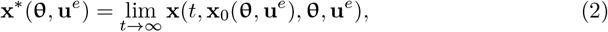

where **x**_0_(**θ, u**^*e*^) is specified in S1 Text, Table S7.

### Observation model

The dependence of observables 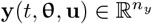 on the state variables and parameters is modeled as

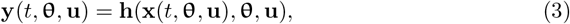

with output map 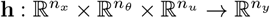. As measurements are corrupted by noise, we model them as noise-corrupted realizations of the observables. The experimental data for these observables are denoted by 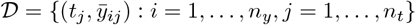, with measurement time points *t*_*j*_ and measurements 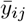. We use common assumptions of measurement independence and additive normally distributed noise:

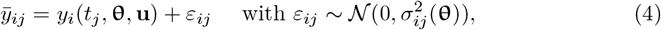

in which *σ*_*ij*_(**θ**) denotes the standard deviation of the noise. More detailed description of the observable is presented in S1 Text, “Observation model” section.

### Parameter estimation

The maximum a posteriori (MAP) estimates of the model parameters are determined by maximizing the posterior probability

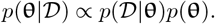

The prior distribution is denoted by *p*(**θ**) and the likelihood is denoted by *p*(𝒟 | **θ**). Here, we assume additive normally distributed measurement noise, yielding the likelihood function

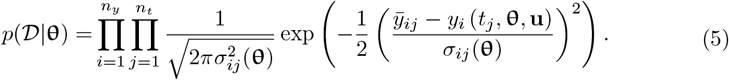

The parameters are assumed to be range bound, and use normally distributed priors,

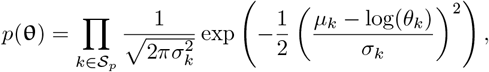

with 𝒮_*p*_ indicating the set of parameters for which prior distributions are available, and *µ*_*k*_ and *σ*_*k*_ indicating the mean and the standard deviation of the prior. The *µ*_*k*_ and *σ*_*k*_ values are given in S1 Text, “Parameter priors”. The parameters **θ** are mostly the logarithms of the reaction rate constants and the observation parameters.

The MAP estimate is obtained by solving the optimization problem

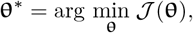

with *J* denoting the negative log posterior,

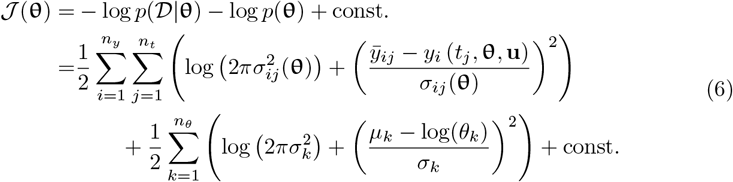

To explore the parameter space, we use a multi-start approach, where multiple local optimizations are performed using randomly sampled starting points [45]. More details on starting point sampling are given in S1 Text, “Parameter starting point sampling”.

In addition, computing the initial conditions required steady-state calculations at each optimization step, increasing the overall computational complexity. To mitigate this, we leveraged a tailored Forward Sensitivity Analysis (FSA) approach, which significantly accelerated model pre-equilibration [51]. As a result, the computation time was reduced by three times compared to using the standard FSA approach (S1 Text, Fig. S2).

### Uncertainty analysis

The parameter credible intervals CI(*θ*_*i*_) were computed from profile posteriors PP(*θ*_*i*_) by

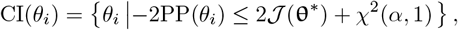

where

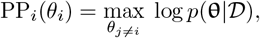

and *χ*^2^(*α*, 1) denotes the *α* quantile of the *χ*^2^-distribution with one degree of freedom. We constructed credible intervals using the 90th, 95th and 99th percentiles [46].

To assess prediction uncertainty, we used an ensemble-based approach. Therefore, we created an ensemble based on all parameter iterates **θ** from optimization that satisfy

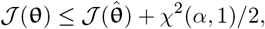

where 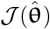 is the smallest objective function value found during optimization. We used the 99th percentile for ensemble construction, i.e. the threshold *χ*^2^(99, 1)*/*2 = 3.32.

## Implementation

The parameter estimation problem was described in the PEtab format [44] and optimization was performed using the Fides gradient-based trust-region optimizer version 0.7.8 [52] via pyPESTO toolbox version 0.5.6 [53]. Model simulations and sensitivity computations were performed using AMICI version 0.32.0 [54].

## Data & code availability

All relevant data for this study are publicly available from the Zenodo repository (https://doi.org/10.5281/zenodo.18619229).

## Supporting information

**S1 Text. Supplementary notes on training data, model details, conserved quantities, observation model, and parameter priors.**(PDF)

## Author contributions

Conceptualization: J.H., J.I., T.H.; Formal Analysis: P.L., J.H.; Funding Acquisition: J.H., T.H. Investigation: P.L., J.I.; Methodology: P.L., J.H., D.W.; Supervision: J.H., D.W., T.H.; Visualization: P.L.; Writing – Original Draft Preparation: P.L., J.H.; Writing – Review & Editing: all authors.

