## Supporting Information for "A mechanistic model of protein kinase A dynamics under pro- and anti-nociceptive inputs"

#### Contents

**S1 Data** **S2**

**S2 Model reactions** **S2**

**S3 Initial values used for pre-equilibration** **S4**

**S4 Conserved quantities** **S4**

**S5 Observation model** **S6**

**S6 Scaling parameters in validation experiments** **S10**

**S7 Parameter priors** **S10**

**S8 Parameter starting point sampling** **S10**

**S9 Supplementary figures** **S12**

### S1 Data

For the estimation of unknown model parameters, we employed a dataset consisting of 12 experimental panels encompassing 162 conditions, including perturbations with forskolin, IBMX, H-89, four cAMP analogs, 5-HT, DAMGO, and fentanyl, resulting in a total of 1,730 data points (Table S1).

### S2 Model reactions

The developed model captures AC activity and AC-mediated cAMP synthesis S2. Forskolin–AC binding and dissociation were described using mass-action kinetics (Table S2, 1 and 2), as was the binding of G-protein $\alpha$ subunits to AC and their subsequent dissociation (Table S2, 3-6). cAMP synthesis is catalyzed by AC, AC:Fsk and AC:$\alpha_S$GTP, reflecting their differential activation states (Table S2, 7-9), whereas cAMP degradation is inhibited by IBMX with inhibition constant $K_{D,IBMX}$ (Table S2, 10).

We assume that cAMP analogs, IBMX, and H-89 are present in excess. Consequently, any consumption of these components through import or binding is considered negligible. Import rates are proportional to the extracellular concentrations of the compounds (Table S2, 11-15).

The model provides a detailed representation of PKA activity. The binding of the PKA catalytic subunit to its inhibitor H-89, as well as its dissociation, was modeled using mass-action kinetics (Table S2, 16,17). cAMP binds to inactive PKA and promotes its activation by inducing dissociation of the regulatory and catalytic subunits (Table S3, 1-4). Similarly to cAMP, Sp-8-Br-cAMPS activates PKA, whereas Rp analogs bind to PKA but do not trigger this dissociation, thereby maintaining the enzyme in its inactive state. To facilitate intracellular delivery, cell-permeable prodrugs of these cAMP analogs were administered to cells; once inside, they are converted into their active forms, which then act on PKA. The RII regulatory subunits of PKA are phosphorylated by the catalytic subunits (Table S3, 5). This phosphorylation reduces their affinity for the catalytic subunit, prolonging PKA activity and modulating the kinetics of holoenzyme reassembly [1]. Accordingly, we account for differing PKA reassembly rates depending on RII phosphorylation state (Table S3, 8 and 9). Binding and dissociation of cAMP and its analogs are modeled according to mass-action kinetics with reaction-specific rate constants (Table S4). It has been demonstrated that RII phosphorylation precedes cAMP binding and controls the inactivation [1], which we take into account in the model.

Ligand binding and dissociation for both MOR and 5-HT$_4$ receptors are modeled according to mass-action kinetics (Table S3, 1-4 and 7-8). We assume that fentanyl, DAMGO, and 5-HT are present in excess, such that their consumption during binding is negligible. Following ligand binding, some receptors may undergo internalization and subsequently be either degraded or recycled to the cell membrane. Because receptor recycling is assumed to be negligible on the timescale considered, it is not included in the model. All mechanisms that render a receptor unavailable are therefore represented collectively as a degradation reaction (Table S3, reactions 5–6 and 9).

The model describes G-protein activation leading to the dissociation of G$\alpha$ and G$\beta\gamma$ complexes, occurring both basally (Table S5, 1 and 3) and in a receptor-mediated manner (Table S5, 2, 4 and 5). It also incorporates G-protein reassembly (Table S5, 6 and 7), as well as GDP–GTP exchange on the G$\alpha$ subunit (Table S5, 8 and 9).

**Table S1.** Summary of experiments used for parameter estimation

| | Experiment Id | Measurement type | Treatments | # |
|---|---|---|---|---|
| datset 1 | JI09_150302_Drg345_343_CycNuc | immunofluorescence microscopy measurements of pRII levels | forskolin, Rp8-Br-cAMPS-pAB, Rp8-pCPT-cAMPS-pAB, Rp-cAMPS-pAB, 4AB-nOH (control) | 144 |
| datset 2 | JI09_150330_Drg350_348_CycNuc | immunofluorescence microscopy measurements of pRII levels | forskolin, IBMX, Sp8-Br-cAMPS-AM | 105 |
| datset 3 | JI09_150330_Drg353_351_CycNuc | immunofluorescence microscopy measurements of pRII levels | forskolin, Rp8-Br-cAMPS-pAB, Rp8-pCPT-cAMPS-pAB, Rp-cAMPS-pAB, 4AB-nOH (control) | 168 |
| datset 4 | JI09_151102_Drg421_418_Age | immunofluorescence microscopy measurements of pRII levels | forskolin, H-89 | 210 |
| datset 5 | JI09_160126_Drg449_444_CycNuc | immunoblotting measurements of pRII | forskolin, IBMX, Sp8-Br-cAMPS-AM | 18 |
| datset 6 | JI09_160201_Drg453_452_CycNuc | immunofluorescence microscopy measurements of C$\alpha$ levels | forskolin, Sp8-Br-cAMPS-AM | 42 |
| datset 7 | LK041_39_MOR_Kinetic_Fentanyl_Fsk | immunofluorescence microscopy measurements of pRII levels | forskolin, fentanyl | 126 |
| datset 8 | LK023_21_MOR_Kinetik_DAMGO_Fsk | immunofluorescence microscopy measurements of pRII levels | forskolin, DAMGO | 126 |
| datset 9 | LK15_150727_LK051_48_MOR _Kinetic_10min_Fentanyl_Fsk | immunofluorescence microscopy measurements of pRII levels | forskolin, fentanyl | 224 |
| datset 10 | LK15_150810_LK053_52_047_46_MOR _Kinetic_Fentanyl_5HT | immunofluorescence microscopy measurements of pRII levels | 5-HT, fentanyl | 119 |
| datset 11 | LK020_18_LK014_12_MOR _Kinetik_DAMGO_5HT | immunofluorescence microscopy measurements of pRII levels | DAMGO, 5-HT | 252 |
| datset 12 | JI09_140331_Drg270_267_TiM | immunofluorescence microscopy measurements of pRII levels | H-89, 5-HT | 196 |

**Table S2.** Model reactions, part I. AC activity and cAMP synthesis, import of IBMX and cAMP analogs.

| # | Reaction | Rate | Note |
|---|---|---|---|
| 1 | AC → AC:Fsk | $k_{f,Fsk}$ [AC][Fsk] | forskolin binding to AC |
| 2 | AC:Fsk → AC | $k_{f,Fsk}$ $K_{D,Fsk}$ [AC:Fsk] | forskolin dissociation from AC |
| 3 | AC + $\alpha_I$GTP → AC:$\alpha_I$GTP | $k_{f,AC:\alpha_I GTP}$ [AC][$\alpha_I$GTP] | AC interaction with G proteins |
| 4 | AC:$\alpha_I$GTP → AC + $\alpha_I$GTP | $k_{f,AC:\alpha_I GTP} K_{D,AC:\alpha_I GTP}$[AC:$\alpha_I$GTP] | |
| 5 | AC + $\alpha_S$GTP → AC:$\alpha_S$GTP | $k_{f,AC:\alpha_S GTP}$[AC][$\alpha_S$GTP] | |
| 6 | AC:$\alpha_S$GTP → AC + $\alpha_S$GtP | $k_{f,AC:\alpha_S GTP} K_{D,AC:\alpha_S GTP}$ [AC:$\alpha_S$GTP] | |
| 7 | ∅ → cAMP | $k_{s,AC,\ cAMP}$ [AC] | cAMP synthesis |
| 8 | ∅ → cAMP | $k_{s,AC,cAMP}$ $\xi_{AC,cAMP,Fsk}$ [AC:Fsk] | |
| 9 | ∅ → cAMP | $k_{s,AC,cAMP}$ $\xi_{AC,cAMP,\alpha_S GTP}$ [AC:$\alpha_S$GTP] | |
| 10 | cAMP → ∅ | $\frac{K_{D,IBMX} \cdot k_{deg,cAMP,free}[cAMP]}{IBMX+K_{D,IBMX}}$ | cAMP degradation |
| 11 | ∅ → IBMX | $k_{i,IBMX}$ ([IBMXex] − [IBMX]) | IBMX import |
| 12 | ∅ → Sp8-Br-cAMPS | $k_{i,Sp8\text{-}Br\text{-}cAMPS\text{-}AM}$ ([Sp8-Br-cAMPS-AM] − [Sp8-Br-cAMPS]) | cAMP analog import |
| 13 | ∅ → Rp8-Br-cAMPS | $k_{i,Rp8\text{-}Br\text{-}cAMPS\text{-}pAB}$ ([Rp8-Br-cAMPS-pAB] − [Rp8-Br-cAMPS]) | |
| 14 | ∅ → Rp8-pCPT-cAMPS | $k_{i,Rp8\text{-}pCPT\text{-}cAMPS\text{-}pAB}$ ([Rp8-pCPT-cAMPS-pAB] − [Rp8-pCPT-cAMPS]) | |
| 15 | ∅ → Rp-cAMPS | $k_{i,Rp\text{-}cAMPS\text{-}pAB}$([Rp-cAMPS-pAB] − [Rp-cAMPS]) | |
| 16 | C → C:H-89 | $k_{f,H\text{-}89}$[C][H-89] | H-89 binding to C |
| 17 | C:H-89 → C | $k_{f,H\text{-}89} K_{D,H\text{-}89}$[C:H-89] | H-89 dissociation from C |

### S3 Initial values used for pre-equilibration

The initial condition for the model is defined as the steady state corresponding to the control experimental condition and is computed by pre-equilibration input. The initial values used for pre-equilibration are listed in Table S7.

### S4 Conserved quantities

In order to take advantage of a more efficient method for sensitivities computation at steady state [2], we reformulated the model by removing conserved quantities. This method requires the system Jacobian to be non-singular, a condition that is violated when conserved quantities are present. The model originally included several conserved quantities:

1. AC + AC:Fsk + AC:$\alpha_I$GTP + AC:$\alpha_S$GTP = const;

**Table S3.** Model reactions, part II. PKA dynamics.

| # | Reaction | Rate | Note |
|---|---|---|---|
| 1 | $pRII_2{:}C_2{:}cAMP_4 \rightarrow C + pRII_2{:}cAMP_4$ | $k_{f,pRII_2:C_2:cAMP_4,pRII_2}\,[pRII_2{:}C_2{:}cAMP_4]$ | PKA activation |
| 2 | $pRII_2{:}C_2{:}Sp8\text{-}Br\text{-}cAMPS_4 \rightarrow C + pRII_2{:}Sp8\text{-}Br\text{-}cAMPS_4$ | $k_{f,pRII_2:C_2:cAMP_4,pRII_2}\,[pRII_2{:}C_2{:}Sp8\text{-}Br\text{-}cAMPS_4]$ | PKA activation |
| 3 | $RII_2{:}C_2 \rightarrow C$ | $k_{f,RII_2:C_2,RII_2}\,[RII_2{:}C_2]$ | PKA activation |
| 4 | $pRII_2{:}C_2 \rightarrow C + pRII_2$ | $k_{f,RII_2:C_2,RII_2}\,\xi_{k_{f,RII_2:C_2,RII_2}}[pRII_2{:}C_2]$ | PKA activation |
| 5 | $RII_2{:}C_2 \rightarrow pRII_2{:}C_2$ | $k_{f,RII_2:C_2,pRII_2:C_2}[RII_2{:}C_2]$ | PKA phosphorylation |
| 6 | $pRII_2{:}C_2 \rightarrow RII_2{:}C_2$ | $k_{f,pRII_2:C_2,RII_2:C_2}[pRII_2{:}C_2]$ | PKA dephosphorylation |
| 7 | $pRII_2 \rightarrow RII_2$ | $k_{f,pRII_2,RII_2}\,[pRII_2]$ | PKA dephosphorylation |
| 8 | $C + RII_2 \rightarrow RII_2{:}C_2$ | $k_{f,RII_2,RII_2:C_2}\,[C][RII_2]$ | PKA deactivation |
| 9 | $C + pRII_2 \rightarrow pRII_2{:}C_2$ | $k_{f,RII_2,RII_2:C_2}\,\xi_{k_{f,RII_2,RII_2:C_2}}[C][pRII_2]$ | PKA deactivation |
| 10 | $pRII_2{:}C_2{:}cAMP_4 \rightarrow pRII_2{:}C_2$ | $\frac{K_{D,IBMX}k_{deg,cAMP}[pRII_2:C_2:cAMP_4]}{IBMX+K_{D,IBMX}}$ | cAMP degradation |
| 11 | $pRII_2{:}cAMP_4 \rightarrow pRII_2$ | $\frac{K_{D,IBMX}k_{deg,cAMP}[pRII_2:cAMP_4]}{IBMX+K_{D,IBMX}}$ | cAMP degradation |

2. $RII_2{:}C_2 + pRII_2{:}C_2 + pRII_2{:}C_2{:}cAMP_4 + pRII_2{:}cAMP_4 + pRII_2{:}C_2{:}Rp8\text{-}Br\text{-}cAMPS_4 + pRII_2{:}C_2{:}Rp8\text{-}pCPT\text{-}cAMPS_4 + pRII_2{:}C_2{:}Rp\text{-}cAMPS_4 + pRII_2{:}C_2{:}Sp8\text{-}Br\text{-}cAMPS_4 + pRII_2{:}Sp8\text{-}Br\text{-}cAMPS_4 + pRII_2 + RII_2 = \text{const}$;

3. $C + C{:}H\text{-}89 + RII_2{:}C_2 + pRII_2{:}C_2 + pRII_2{:}C_2{:}cAMP_4 + pRII_2{:}C_2{:}Rp8\text{-}Br\text{-}cAMPS_4 + pRII_2{:}C_2{:}Rp8\text{-}pCPT\text{-}cAMPS_4 + pRII_2{:}C_2{:}Rp\text{-}cAMPS_4 + pRII_2{:}C_2{:}Sp8\text{-}Br\text{-}cAMPS_4 = \text{const}$;

4. $\alpha_I\beta_I\gamma_I + \alpha_I GTP + \alpha_I GDP + AC{:}\alpha_I GTP = \text{const}$;

5. $\alpha_S\beta_S\gamma_S + \alpha_S GTP + \alpha_S GDP + AC{:}\alpha_S GTP = \text{const}$;

6. $\alpha_I\beta_I\gamma_I + \beta_I\gamma_I = \text{const}$;

7. $\alpha_S\beta_S\gamma_S + \beta_S\gamma_S = \text{const}$.

The conserved quantities can be removed by solving each equation for one of the state variables, and thereby reducing the model dimension by 7. The following 7 state variables were removed from the model: AC:Fsk, $RII_2$, C:H-89, $\alpha_I GDP$, $\alpha_S GDP$, $\beta_I\gamma_I$ and $\beta_S\gamma_S$.

Depending on the experimental condition, certain parts of the network may be included or excluded. In the absence of opioids, additional conserved quantities emerge due to the absence of reactions involving ligands. This is the case in the control condition used to compute the system's initial state. To address this, we introduced receptor degradation reactions that are active only when receptor ligands are absent. These reactions do not influence downstream dynamics, ensuring that they do not affect the system's behavior under other conditions.

**Table S4.** Model reactions, part II. Binding and dissociation of cAMP and its analogs.

| # | Reaction | Rate | Note |
|---|---|---|---|
| 1 | $pRII_2$:$C_2$ → $pRII_2$:$C_2$:$cAMP_4$ | $k_{f,cAMP}$ [$pRII_2$:$C_2$][cAMP] | cAMP binding |
| 2 | $pRII_2$:$C_2$:$cAMP_4$ → $pRII_2$:$C_2$ | $k_{f,cAMP}$ $K_{D,cAMP}$ [$pRII_2$:$C_2$:$cAMP_4$] | cAMP dissociation from inactive PKA |
| 3 | $pRII_2$:$cAMP_4$ → $pRII_2$ | $k_{f,cAMP}$ $K_{D,cAMP}$[$pRII_2$:$cAMP_4$] | cAMP dissociation from regulatory subunit |
| 4 | $pRII_2$:$C_2$ → $pRII_2$:$C_2$:Rp8-Br-$cAMPS_4$ | $k_{f,cAMP}$ $\xi_{b,Rp8\text{-}Br\text{-}cAMPS}$ [$pRII_2$:$C_2$][Rp8-Br-cAMPS] | cAMP analog binding to PKA |
| 5 | $pRII_2$:$C_2$:Rp8-Br-$cAMPS_4$ → $pRII_2$:$C_2$ | $k_{f,cAMP}$ $K_{D,cAMP}$ $\xi_{b,Rp8\text{-}Br\text{-}cAMPS}$ $\xi_{K_D,Rp8\text{-}Br\text{-}cAMPS}$ [$pRII_2$:$C_2$:Rp8-Br-$cAMPS_4$] | cAMP analog dissociation |
| 6 | $pRII_2$:$C_2$ → $pRII_2$:$C_2$:Rp8-pCPT-$cAMPS_4$ | $k_{f,cAMP}$ $\xi_{b,Rp8\text{-}pCPT\text{-}cAMPS}$ [$pRII_2$:$C_2$][Rp8-pCPT-cAMPS] | cAMP analog binding to PKA |
| 7 | $pRII_2$:$C_2$:Rp8-pCPT-$cAMPS_4$ → $pRII_2$:$C_2$ | $k_{f,cAMP}$ $K_{D,cAMP}$ $\xi_{b,Rp8\text{-}pCPT\text{-}cAMPS}$ $\xi_{K_D,Rp8\text{-}pCPT\text{-}cAMPS}$ [$pRII_2$:$C_2$:Rp8-pCPT-$cAMPS_4$] | cAMP analog dissociation |
| 8 | $pRII_2$:$C_2$ → $pRII_2$:$C_2$:Rp-$cAMPS_4$ | $k_{f,cAMP}$ $\xi_{b,Rp\text{-}cAMPS}$ [$pRII_2$:$C_2$][Rp-cAMPS] | cAMP analog binding to PKA |
| 9 | $pRII_2$:$C_2$:Rp-$cAMPS_4$ → RIIp_C_2 | $k_{f,cAMP}$ $K_{D,cAMP}$ $\xi_{b,Rp\text{-}cAMPS}$ $\xi_{K_D,Rp\text{-}cAMPS}$ [$pRII_2$:$C_2$:Rp-$cAMPS_4$] | cAMP analog dissociation |
| 10 | $pRII_2$:$C_2$ → $pRII_2$:$C_2$:Sp8-Br-$cAMPS_4$ | $k_{f,cAMP}$ $\xi_{b,Sp8\text{-}Br\text{-}cAMPS}$ [$pRII_2$:$C_2$] [Sp8-Br-cAMPS] | cAMP analog binding to PKA |
| 11 | $pRII_2$:$C_2$:Sp8-Br-$cAMPS_4$ → $pRII_2$:$C_2$ | $k_{f,cAMP}$ $K_{D,cAMP}$ $\xi_{b,Sp8\text{-}Br\text{-}cAMPS}$ $\xi_{K_D,Sp8\text{-}Br\text{-}cAMPS}$ [$pRII_2$:$C_2$:Sp8-Br-$cAMPS_4$] | cAMP analog dissociation |
| 12 | $pRII_2$:Sp8-Br-$cAMPS_4$ → $pRII_2$ | $k_{f,cAMP}$ $K_{D,cAMP}$ $\xi_{b,Sp8\text{-}Br\text{-}cAMPS}$ $\xi_{K_D,Sp8\text{-}Br\text{-}cAMPS}$ [$pRII_2$:Sp8-Br-$cAMPS_4$] | cAMP dissociation |

### S5 Observation model

There are three observables in the available data: signal intensities of pRII and $C_\alpha$ from microscopy images, and Western blot measurements of pRII. Assuming normally distributed measurement noise with unknown standard deviations, we define the following observation model:

1. For the total pRII abundance determined by microscopy measurements we use

$$y_{i,j} = s_j \cdot (s_{\text{pRII,global}} \cdot \text{pRII}^i_{\text{total}} + b_{\text{pRII,global}} \cdot b_j) + \varepsilon_i, \quad \varepsilon_i \sim \mathcal{N}(0, \sigma_j). \tag{1}$$

   Here, $j \in \{1, \ldots, n_e\}$, $n_e$ is the number of experiments, $i \in \{1, \ldots, n_j\}$, $n_j$ is the number of measurements corresponding to the experiment $j$, $s_{\text{pRII,global}}$ is the scaling parameter and $b_{\text{pRII,global}}$ is the offset parameter for all pRII microscopy measurements, $s_j$ and $b_j$ are the experiment-specific scaling and offset parameters, respectively. Data analysis revealed that the offset parameter $b_j$ can be set to one

**Table S5.** Model reactions, part III. Receptor dynamics.

| # | Reaction | Rate | Note |
|---|---|---|---|
| 1 | MOR → MOR:DAMGO | $k_{f,DAMGO}$ [MOR][DAMGO] | DAMGO binding to MOR |
| 2 | MOR:DAMGO → MOR | $K_{D,DAMGO}k_{f,DAMGO}$ [MOR:DAMGO] | DAMGO dissociation from MOR |
| 3 | MOR → MOR:Fentanyl | $k_{f,Fentanyl}$ [MOR][Fentanyl] | Fentanyl binding to MOR |
| 4 | MOR:Fentanyl → MOR | $k_{f,Fentanyl}K_{D,Fentanyl}$ [MOR:Fentanyl] | Fentanyl dissociation from MOR |
| 5 | MOR:DAMGO → ∅ | $k_{int,DAMGO}$ [MOR:DAMGO] | receptor internalization |
| 6 | MOR:Fentanyl → ∅ | $k_{int,Fentanyl}$ [MOR:Fentanyl] | receptor internalization |
| 7 | 5-HT$_4$ → 5-HT$_4$:5-HT | $k_{f,5\text{-}HT}$ [5-HT$_4$][5-HT] | 5-HT binding to 5-HT$_4$ |
| 8 | 5-HT$_4$:5-HT → 5-HT4 | $K_{D,5\text{-}HT}$ $k_{f,5\text{-}HT}$ [5-HT$_4$:5-HT] | 5-HT dissociation from 5-HT$_4$ |
| 9 | 5-HT$_4$:5-HT → ∅ | $k_{int,5\text{-}HT}$ [5-HT$_4$:5-HT] | receptor internalization |

for all experiments except one, LK15_150727_LK051_48_MOR_Kinetic_10min_Fentanyl_Fsk, for which it was estimated from the data. The experiment-specific noise parameters are defined by $\sigma_j = s_j \cdot s_{\mathrm{pRII,global}} \cdot \rho_{\mathrm{pRII,Microscopy}}$, where $\rho_{\mathrm{pRII,Microscopy}}$ is the noise parameter that need to be estimated from the data, and

$$\begin{aligned}\mathrm{pRII}^i_{\mathrm{total}} =& (2 \cdot \mathrm{pRII}^i_2 + 2 \cdot \mathrm{pRII_2{:}cAMP}^i_4 + 2 \cdot \mathrm{pRII_2{:}Sp8\text{-}Br\text{-}cAMPS}^i_4 \\ &+2 \cdot \mathrm{rel_{open}}(\mathrm{pRII_2{:}C}^i_2 + \mathrm{pRII_2{:}C_2{:}Rp\text{-}cAMPS}^i_4 \\ &\quad + \mathrm{pRII_2{:}C_2{:}Rp8\text{-}Br\text{-}cAMPS}^i_4 + \mathrm{pRII_2{:}C_2{:}Rp8\text{-}pCPT\text{-}cAMPS}^i_4) \\ &+2 \cdot (\mathrm{rel_{open}} - \xi_{\mathrm{rel_{open}}}(\mathrm{rel_{open}} - 1))(\mathrm{pRII_2{:}C_2{:}cAMP}^i_4 \\ &\quad + \mathrm{pRII_2{:}C_2{:}Sp8\text{-}Br\text{-}cAMPS}^i_4)),\end{aligned}$$

with $\mathrm{rel_{open}} \in [10^{-5}, 1]$, $\xi_{\mathrm{rel_{open}}} \in [10^{-5}, 1]$, (Fig. S1a). This formula takes into account that complexes with pRII have different antibody accessibility, depending on whether the C subunit and cAMP or its analogs are bound to them.

2. Similarly to pRII, for the $C_\alpha$ microscopy the measurements observation model is

$$y_i = s_{C_\alpha} \cdot (s_{C_\alpha,\mathrm{global}} \cdot C^i_{\alpha,\mathrm{total}} + b_{C_\alpha,\mathrm{global}}) + \varepsilon_{C_\alpha,i}, \quad \varepsilon_{C_\alpha,i} \sim \mathcal{N}(0, \sigma_{C_\alpha}), \tag{2}$$

where $\sigma_{C_\alpha} = s_{C_\alpha} \cdot s_{C_\alpha,\mathrm{global}} \cdot \rho_{C_\alpha,\mathrm{Microscopy}}$ and

$$\begin{aligned}C^i_{\alpha,\mathrm{total}} =& 2 \cdot \mathrm{C} + 2 \cdot \mathrm{C{:}H\text{-}89} \\ &+2 \cdot \mathrm{rel_{open}}(\mathrm{pRII_2{:}C_2} + \mathrm{pRII_2{:}C_2{:}Rp\text{-}cAMPS_4} \\ &\quad + \mathrm{pRII_2{:}C_2{:}Rp8\text{-}Br\text{-}cAMPS_4} + \mathrm{pRII_2{:}C_2{:}Rp8\text{-}pCPT\text{-}cAMPS_4}) \\ &+2 \cdot (\mathrm{rel_{open}} - \xi_{\mathrm{rel_{open}}}(\mathrm{rel_{open}} - 1))(\mathrm{pRII_2{:}C_2{:}cAMP_4} \\ &\quad + \mathrm{pRII_2{:}C_2{:}Sp8\text{-}Br\text{-}cAMPS_4}),\end{aligned}$$

**Table S6.** Model reactions, part IV. G protein dynamics.

| # | Reaction | Rate | Note |
|---|---|---|---|
| 1 | $\alpha_S\beta_S\gamma_S \rightarrow \alpha_S GTP + \beta_S\gamma_S$ | $k_{f,\alpha_S\beta_S\gamma_S}\ [\alpha_S\beta_S\gamma_S]$ | $G_S$ protein activation |
| 2 | $\alpha_S\beta_S\gamma_S \rightarrow \alpha_S GTP + \beta_S\gamma_S$ | $k_{f,\alpha_S\beta_S\gamma_S}\ \xi_{\alpha_S\beta_S\gamma_S,5\text{-HT}_4,5\text{-HT}}\ [5\text{-HT}_4{:}5\text{-HT}]\ [\alpha_S\beta_S\gamma_S]$ | $G_S$ protein activation mediated by 5-HT$_4$ |
| 3 | $\alpha_I\beta_I\gamma_I \rightarrow \alpha_I GTP + \beta_I\gamma_I$ | $k_{f,\alpha_I\beta_I\gamma_I}\ [\alpha_I\beta_I\gamma_I]$ | $G_I$ protein activation |
| 4 | $\alpha_I\beta_I\gamma_I \rightarrow \alpha_I GTP + \beta_I\gamma_I$ | $k_{f,\alpha_I\beta_I\gamma_I}\ \xi_{\alpha_I\beta_I\gamma_I,\text{MOR},\text{DAMGO}}\ [\text{MOR:DAMGO}][\alpha_I\beta_I\gamma_I]$ | $G_I$ protein activation mediated my MORs |
| 5 | $\alpha_I\beta_I\gamma_I \rightarrow \alpha_I GTP + \beta_I\gamma_I$ | $k_{f,\alpha_I\beta_I\gamma_I}\ \xi_{\alpha_I\beta_I\gamma_I,\text{MOR},\text{Fentanyl}}\ [\text{MOR:Fentanyl}]\ [\alpha_I\beta_I\gamma_I]$ | $G_I$ protein activation mediated my MORs |
| 6 | $\alpha_I GDP + \beta_I\gamma_I \rightarrow \alpha_I\beta_I\gamma_I$ | $k_{f,\alpha_I GDP,\beta_I\gamma_I,\alpha_I\beta_I\gamma_I}\ [\alpha_I GDP][\beta_I\gamma_I]$ | $G_I$ protein reassembly |
| 7 | $\alpha_S GDP + \beta_S\gamma_S \rightarrow \alpha_S\beta_S\gamma_S$ | $k_{f,\alpha_S GDP,\beta_S\gamma_S,\alpha_S\beta_S\gamma_S}\ [\alpha_S GDP][\beta_S\gamma_S]$ | $G_S$ protein reassembly |
| 8 | $\alpha_S GTP \rightarrow \alpha_S GDP$ | $k_{f,\alpha_S GTP,\alpha_S GDP}\ [\alpha_S GTP]$ | hydrolysis of GTP bound to $G_{\alpha_S}$ to GDP |
| 9 | $\alpha_I GTP \rightarrow \alpha_I GDP$ | $k_{f,\alpha_I GTP,\alpha_I GDP}\ [\alpha_I GTP]$ | hydrolysis of GTP bound to $G_{\alpha_I}$ to GDP |

(Fig. S1b). Similarly to $\text{pRII}_i^{\text{total}}$, parameters $\text{rel}_{\text{open}}$ and $\xi_{\text{rel}_{\text{open}}}$ reflect difference in accessibility of the epitope and, therefore, different contribution to the signal intensity. Only one experiment was used for parameter estimation, where $C_\alpha$ was measured, therefore, experiment specific scaling parameters are not needed here.

3. There is only one Western blot experiment and the observation model is

$$y_i = s_{\text{pRII,Western}} \cdot \text{pRII}_i^{\text{Western,total}} + \varepsilon_{\text{Western},i}, \quad \varepsilon_{\text{Western},i} \sim \mathcal{N}(0, \sigma_{\text{pRII,Western}}), \tag{3}$$

where

$$\begin{aligned}\text{pRII}_{\text{Western,total}} &= 2\cdot\text{pRII}_2 + 2\cdot\text{pRII}_2{:}\text{C}_2 + 2\cdot\text{pRII}_2{:}\text{cAMP}_4 + 2\cdot\text{pRII}_2{:}\text{C}_2{:}\text{cAMP}_4\\ &+ 2\cdot\text{pRII}_2{:}\text{C}_2{:}\text{Rp-cAMPS}_4 + 2\cdot\text{pRII}_2{:}\text{Sp8-Br-cAMPS}_4\\ &+ 2\cdot\text{pRII}_2{:}\text{C}_2{:}\text{Rp8-Br-cAMPS}_4 + 2\cdot\text{pRII}_2{:}\text{C}_2{:}\text{Sp8-Br-cAMPS}_4\\ &+ 2\cdot\text{pRII}_2{:}\text{C}_2{:}\text{Rp8-pCPT-cAMPS}_4,\end{aligned}$$

with $\sigma_{\text{pRII,Western}} = s_{\text{pRII,Western}} \cdot \rho_{\text{pRII,Western}}$ (Fig. S1c).

Parameters $s_j$, $s_{\text{pRII,global}}$, $b_{\text{pRII,global}}$, $b_{\text{LK15\_150727\_LK051\_48\_MOR\_Kinetic\_10min\_Fentanyl\_Fsk}}$, $\rho_{\text{pRII,Microscopy}}$, $s_{C_\alpha}$, $s_{C_\alpha,\text{global}}$, $b_{C_\alpha,\text{global}}$, $\rho_{C_\alpha,\text{Microscopy}}$, $s_{\text{pRII,Western}}$, $\rho_{\text{pRII,Western}}$ need to be estimated from the available measurements.

**Table S7.** Initial values used for pre-equilibration using the control experimental condition.

| **Species** | **Initial value** |
|---|---|
| AC | $AC_{total}$ |
| cAMP | 0 |
| Csub | 0 |
| $RII_2$:$C_2$ | 0 |
| $pRII_2$:$C_2$ | RIIp_C_$2_{total}$ |
| $pRII_2$:$C_2$:$cAMP_4$ | 0 |
| $pRII_2$:$cAMP_4$ | 0 |
| Sp8-Br-cAMPS | 0 |
| Rp8-Br-cAMPS | 0 |
| Rp8-pCPT-cAMPS | 0 |
| Rp-cAMPS | 0 |
| $pRII_2$:$C_2$:Rp8-Br-$cAMPS_4$ | 0 |
| $pRII_2$:$C_2$:Rp8-pCPT-$cAMPS_4$ | 0 |
| $pRII_2$:$C_2$:Rp-$cAMPS_4$ | 0 |
| $pRII_2$:$C_2$:Sp8-Br-$cAMPS_4$ | 0 |
| $pRII_2$:Sp8-Br-$cAMPS_4$ | 0 |
| $pRII_2$ | 0 |
| IBMX | 0 |
| 5-$HT_4$ | $fiveHT4_{total}$ |
| 5-$HT_4$:5-HT | 0 |
| MOR | $MOR_{total}$ |
| MOR:DAMGO | 0 |
| MOR:Fentanyl | 0 |
| $\alpha_S\beta_S\gamma_S$ | alphaS_betaS_$gammaS_{total}$ |
| $\alpha_I\beta_I\gamma_I$ | alphaI_betaI_$gammaI_{total}$ |
| $\alpha_S$GTP | 0 |
| $\alpha_I$GTP | 0 |
| AC:$\alpha_S$GTP | 0 |
| AC:$\alpha_I$GTP | 0 |

#### S6 Scaling parameters in validation experiments

In the validation experiments, the observable for pRII was computed according to formula (1). The experiment-specific scaling parameter for dataset 2 was taken from the MAP estimate, since part of this dataset was included in the training data. For datasets 13 and 14, the scaling parameters were estimated by minimizing the discrepancy between the model simulations and the measurements across all experimental conditions within each dataset.

#### S7 Parameter priors

To facilitate parameter estimation and ensure more biologically plausible parameter estimates we specified prior distributions for some estimated parameters (Table S8). We used log-normal prior distributions for dissociation constants $K_{D,Fsk}$, $K_{D,H\text{-}89}$ and $K_{D,cAMP}$, with mean values taken from [3], [4] and [5], respectively. Similarly, log-normal priors were used for parameters $\xi_{k_f,RII_2,RII_2:C_2}$ and $\xi_{k_f,RII_2:C_2,RII_2}$ with mean values taken from [6]. Weakly informative priors were used for parameters that define cAMP analog-specific binding and dissociation rates. Namely, we chose log-normal prior with mean equal to 0 and standard deviation equal to 3. Additionally, we used log-normal priors for the experiment-specific scaling parameters with mean equal to 0 and standard deviation equal to 0.1.

#### S8 Parameter starting point sampling

Due to the high dimensionality of the parameter space, parameter estimation posed a significant challenge. To address this, we leveraged our previous results from [1], using them to inform the sampling of starting points for the 46 parameters estimated in that study. Specifically, the starting points for these parameters were drawn from log-normal distributions with mean values equal to logarithm to base 10 of the best-fit values obtained in [1], and standard deviations set to the absolute values of 20% of the respective means (Tables S9 and S10). The remaining 30 parameters were initialized using log-uniform sampling within predefined bounds.

**Table S8.** Prior information for the parameter estimation.

| **Parameter** | **Mean** | **Std** | **Ref** |
|---|---|---|---|
| model parameters | | | |
| $K_{D,Fsk}$ | 0.8451 | 0.2 | [3] |
| $K_{D,H\text{-}89}$ | -1.3188 | 0.2 | [4] |
| $K_{D,IBMX}$ | 1 | 0.2 | |
| $K_{D,cAMP}$ | 0.4624 | 0.2 | [5] |
| $\xi_{K_D,Rp8\text{-}Br\text{-}cAMPS}$ | 0 | 3 | |
| $\xi_{K_D,Rp8\text{-}pCPT\text{-}cAMPS}$ | 0 | 3 | |
| $\xi_{K_D,Rp\text{-}cAMPS}$ | 0 | 3 | |
| $\xi_{K_D,Sp8\text{-}Br\text{-}cAMPS}$ | 0 | 3 | |
| $\xi_{b,Rp8\text{-}Br\text{-}cAMPS}$ | 0 | 3 | |
| $\xi_{b,Rp8\text{-}pCPT\text{-}cAMPS}$ | 0 | 3 | |
| $\xi_{b,Rp\text{-}cAMPS}$ | 0 | 3 | |
| $\xi_{b,Sp8\text{-}Br\text{-}cAMPS}$ | 0 | 3 | |
| $\xi_{k_f,RII_2,RII_2:C_2}$ | -1.7424 | 0.1 | [6] |
| $\xi_{k_f,RII_2:C_2,RII_2}$ | -0.0621 | 0.1 | [6] |
| observable and noise parameters | | | |
| $b_{C\alpha,global}$ | 2 | 0.1 | |
| $b_{pRII,global}$ | 2 | 0.1 | |
| $s_{pRII,JI09\_150302\_Drg345\_343\_CycNuc}$ | 0 | 0.1 | |
| $s_{pRII,JI09\_150330\_Drg350\_348\_CycNuc}$ | 0 | 0.1 | |
| $s_{pRII,JI09\_150330\_Drg353\_351\_CycNuc}$ | 0 | 0.1 | |
| $s_{pRII,JI09\_151102\_Drg421\_418\_Age}$ | 0 | 0.1 | |
| $s_{pRII,LK041\_39\_MOR\_Kinetic\_Fentanyl\_Fsk}$ | 0 | 0.1 | |
| $s_{pRII,LK023\_21\_MOR\_Kinetik\_DAMGO\_Fsk}$ | 0 | 0.1 | |
| $s_{pRII,LK15\_150810\_LK053\_52\_047\_46\_MOR\_Kinetic\_Fentanyl\_5HT}$ | 0 | 0.1 | |
| $s_{pRII,LK15\_150727\_LK051\_48\_MOR\_Kinetic\_10min\_Fentanyl\_Fsk}$ | 0 | 0.1 | |
| $s_{pRII,LK020\_18\_LK014\_12\_MOR\_Kinetik\_DAMGO\_5HT}$ | 0 | 0.1 | |
| $s_{pRII,JI09\_140331\_Drg270\_267\_TiM}$ | 0 | 0.1 | |

**Table S9.** Sampling distribution parameters used for sampling starting points for parameter estimation, model parameters.

| Parameter | Mean | Std |
|---|---|---|
| $K_{D,Fsk}$ | 1.15 | 0.231 |
| $K_{D,H\text{-}89}$ | -1.32 | 0.264 |
| $K_{D,IBMX}$ | 1.08 | 0.216 |
| $K_{D,cAMP}$ | 0.45 | 0.091 |
| $k_{deg,cAMP}$ | -5.0 | 1.0 |
| $k_{deg,cAMP,free}$ | 0.87 | 0.174 |
| $k_{f,Fsk}$ | 3.0 | 0.6 |
| $k_{f,H\text{-}89}$ | -3.13 | 0.626 |
| $k_{f,RII_2,RII_2:C_2}$ | 0.1 | 0.02 |
| $k_{f,RII_2:C_2,RII_2}$ | -1.73 | 0.345 |
| $k_{f,RII_2:C_2,pRII_2:C_2}$ | -1.64 | 0.329 |
| $k_{f,pRII_2,RII_2}$ | -1.49 | 0.298 |
| $k_{f,pRII_2:C_2,RII_2:C_2}$ | -1.59 | 0.318 |
| $k_{f,pRII_2:C_2:cAMP_4,pRII_2}$ | -0.98 | 0.196 |
| $k_{f,cAMP}$ | -0.3 | 0.06 |
| $k_{i,IBMX}$ | 3.0 | 0.6 |
| $k_{i,Rp8\text{-}Br\text{-}cAMPS\text{-}pAB}$ | 0.81 | 0.162 |
| $k_{i,Rp8\text{-}pCPT\text{-}cAMPS\text{-}pAB}$ | 0.7 | 0.195 |
| $k_{i,Rp\text{-}cAMPS\text{-}pAB}$ | -1.79 | 0.357 |
| $k_{i,Sp8\text{-}Br\text{-}cAMPS\text{-}AM}$ | -0.88 | 0.176 |
| $k_{s,AC,cAMP}$ | -0.37 | 0.075 |
| $\xi_{AC,cAMP,Fsk}$ | 2.81 | 0.562 |
| $\xi_{K_D,Rp8\text{-}Br\text{-}cAMPS}$ | -1.4 | 0.28 |
| $\xi_{K_D,Rp8\text{-}pCPT\text{-}cAMPS}$ | -0.71 | 0.142 |
| $\xi_{K_D,Rp\text{-}cAMPS}$ | -0.79 | 0.158 |
| $\xi_{K_D,Sp8\text{-}Br\text{-}cAMPS}$ | -0.65 | 0.13 |
| $\xi_{b,Rp8\text{-}Br\text{-}cAMPS}$ | -1.39 | 0.278 |
| $\xi_{b,Rp8\text{-}pCPT\text{-}cAMPS}$ | -1.57 | 0.315 |
| $\xi_{b,Rp\text{-}cAMPS}$ | -0.22 | 0.044 |
| $\xi_{b,Sp8\text{-}Br\text{-}cAMPS}$ | 1.33 | 0.267 |
| $\xi_{k_f,RII_2,RII_2:C_2}$ | -1.72 | 0.345 |
| $\xi_{k_f,RII_2:C_2,RII_2}$ | -0.09 | 0.018 |

### S9 Supplementary figures

2. Lakrisenko P, Pathirana D, Weindl D, Hasenauer J. Benchmarking methods for computing local

**Table S10.** Sampling distribution parameters used for sampling starting points for parameter estimation, observable and noise parameters.

| Parameter | Mean | Std |
|---|---|---|
| $b_{C\alpha,global}$ | 2.01 | 0.401 |
| $b_{pRII,global}$ | 2.0 | 0.4 |
| $rel_{open}$ | -0.63 | 0.127 |
| $s_{C\alpha,global}$ | 3.25 | 0.649 |
| $s_{pRII,JI09\_150302\_Drg345\_343\_CycNuc}$ | 0.04 | 0.009 |
| $s_{pRII,JI09\_150330\_Drg350\_348\_CycNuc}$ | -0.04 | 0.009 |
| $s_{pRII,JI09\_150330\_Drg353\_351\_CycNuc}$ | -0.03 | 0.006 |
| $s_{pRII,JI09\_151102\_Drg421\_418\_Age}$ | 0.03 | 0.006 |
| $s_{pRII,Western}$ | 0.06 | 0.012 |
| $s_{pRII,global}$ | 3.34 | 0.669 |
| $\rho_{C\alpha,Microscopy}$ | -1.3 | 0.261 |
| $\rho_{pRII,Microscopy}$ | -1.39 | 0.278 |
| $\rho_{pRII,Western}$ | -1.0 | 0.201 |
| $\xi_{rel_{open}}$ | -0.13 | 0.025 |


sensitivities in ordinary differential equation models at dynamic and steady states. PLOS ONE. 2024;19(10):1–19. doi:10.1371/journal.pone.0312148.

3. Awad JA, Johnson RA, Jakobs KH, Schultz G. Interactions of forskolin and adenylate cyclase. Effects on substrate kinetics and protection against inactivation by heat and N-ethylmaleimide. J Biol Chem. 1983;258(5):2960–2965.

4. Lochner A, Moolman JA. The many faces of H89: a review. Cardiovasc Drug Rev. 2006;24(3-4):261–274.

5. Dao KK, Teigen K, Kopperud R, Hodneland E, Schwede F, Christensen AE, et al. Epac1 and cAMP-dependent protein kinase holoenzyme have similar cAMP affinity, but their cAMP domains have distinct structural features and cyclic nucleotide recognition. J Biol Chem. 2006;281(30):21500–21511.

6. Zhang P, Knape MJ, Ahuja LG, Keshwani MM, King CC, Sastri M, et al. Single Turnover Autophosphorylation Cycle of the PKA RII$\beta$ Holoenzyme. PLOS Biol. 2015;13(7):1–22. doi:10.1371/journal.pbio.1002192.

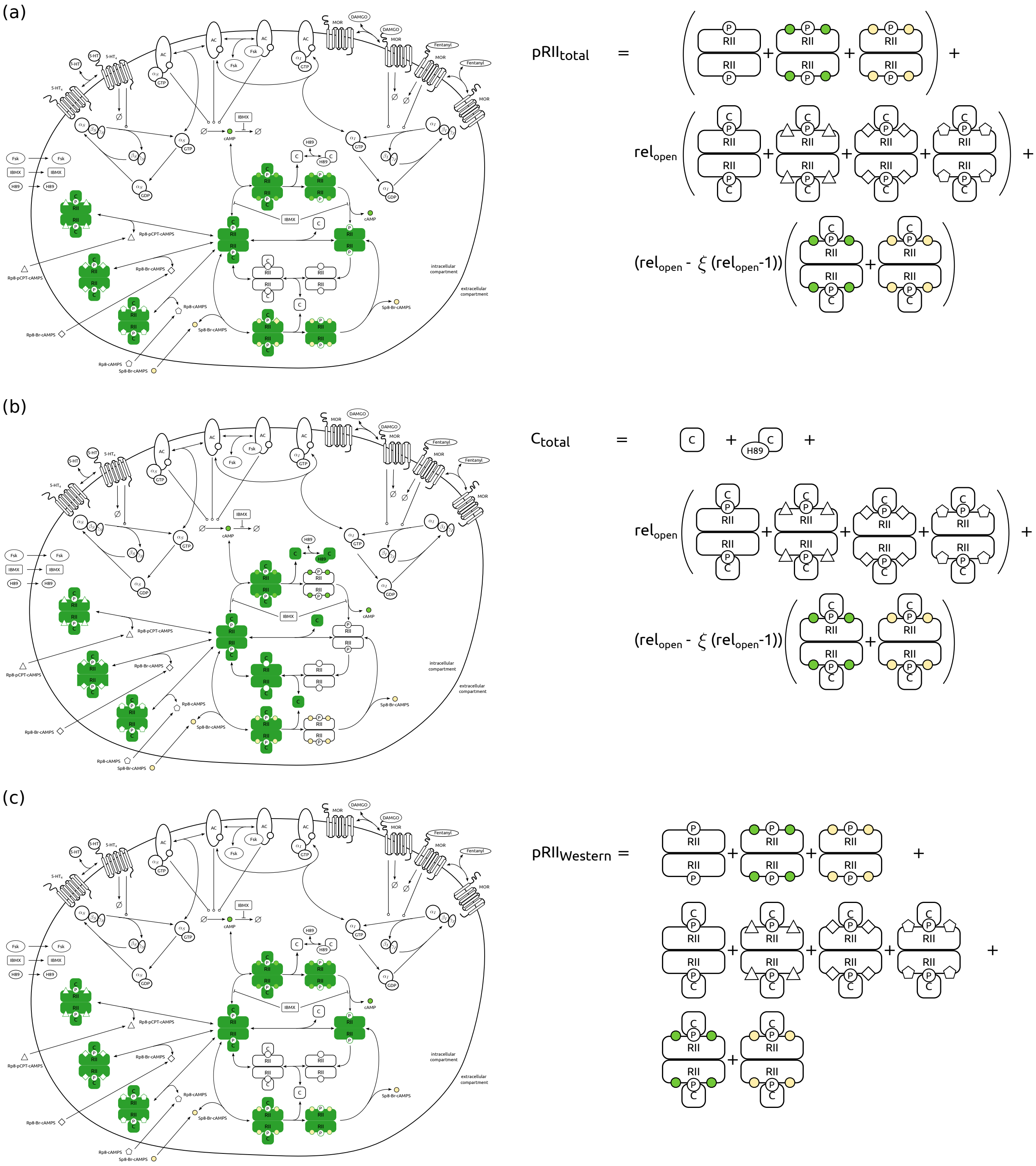


**Fig S1. Observables accessible by the employed measurements techniques.** The green color highlights the species comprising the observables. (a) The observable corresponding to the immunofluorescence microscopy measurements of pRII (b) and $C_\alpha$. (c) The observable corresponding to the Western blot measurements.

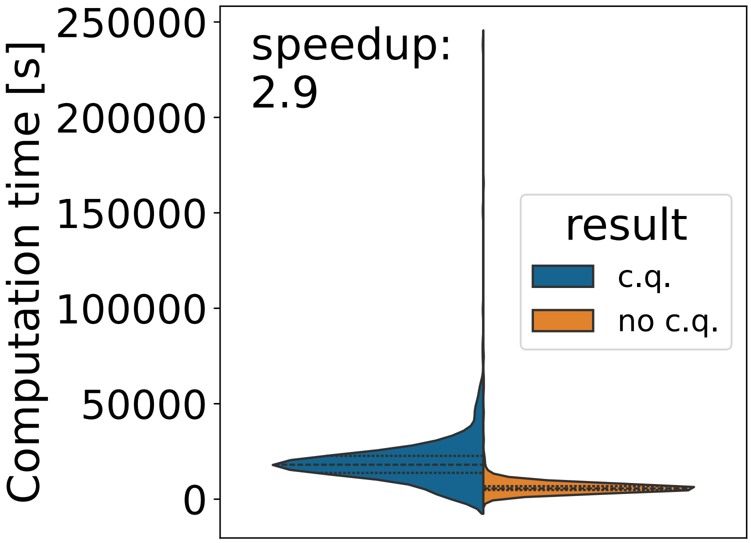


**Fig S2. Optimization computation time speedup from using ssFSA compared to standard FSA approach.**

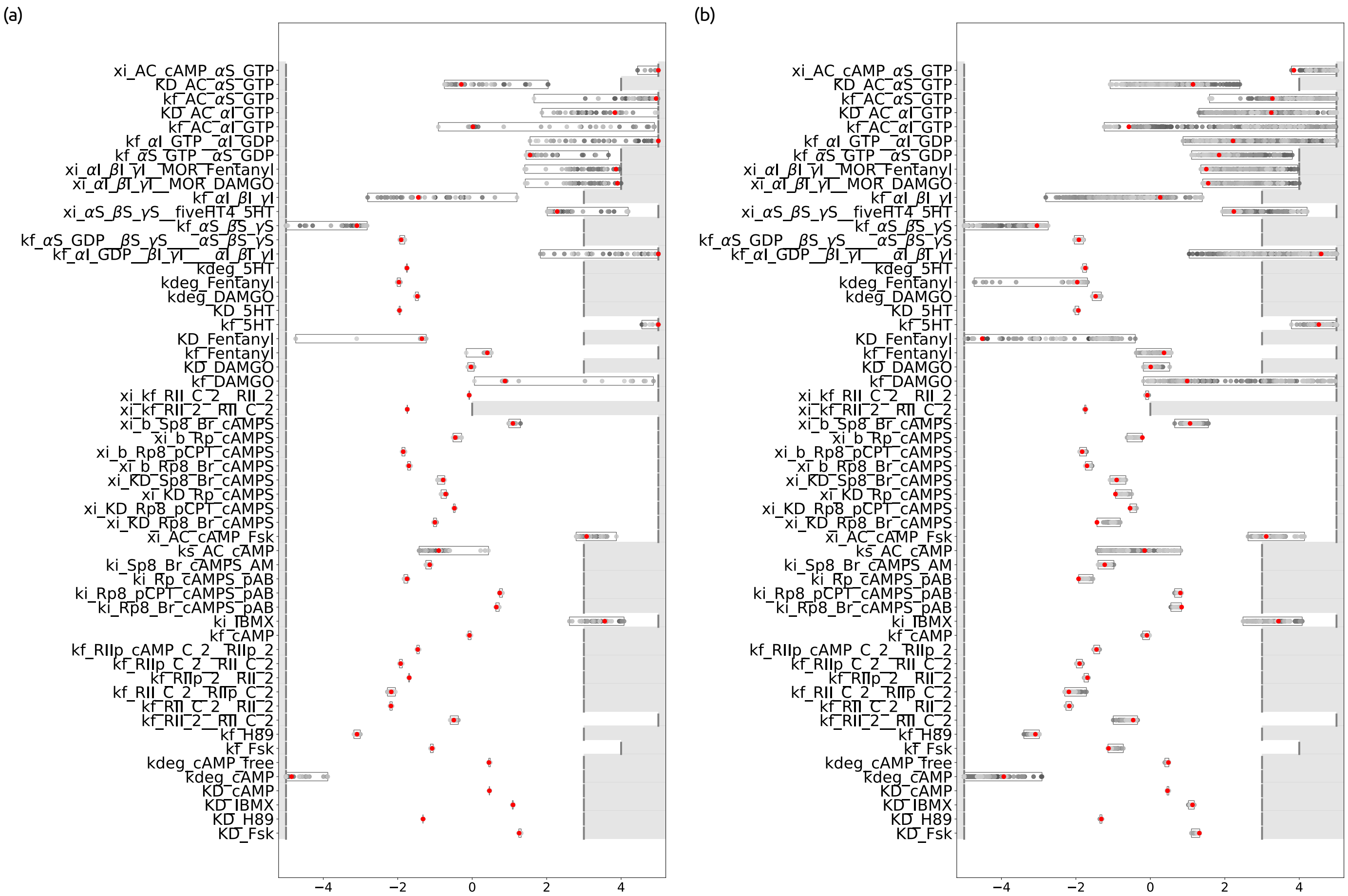


**Fig S3. Parameter ensembles assembled from the parameter vectors encountered during the parameter optimization.** Only model parameters are shown. (a) Ensemble of 37 parameter vectors corresponding to the 37 lowest objective function values; these fits are not statistically distinguishable based on a $\chi^2$ test. (b) Ensemble of all parameter vectors encountered during optimization that yield fits statistically indistinguishable from the maximum a posteriori estimate at a significance level of 0.01, corresponding to an objective function difference of 3.32. Each point represents a parameter value; red points indicate values from the best optimization run. Boxes show the range spanned by parameters in the ensemble, and the white region indicates the parameter bounds used for estimation.

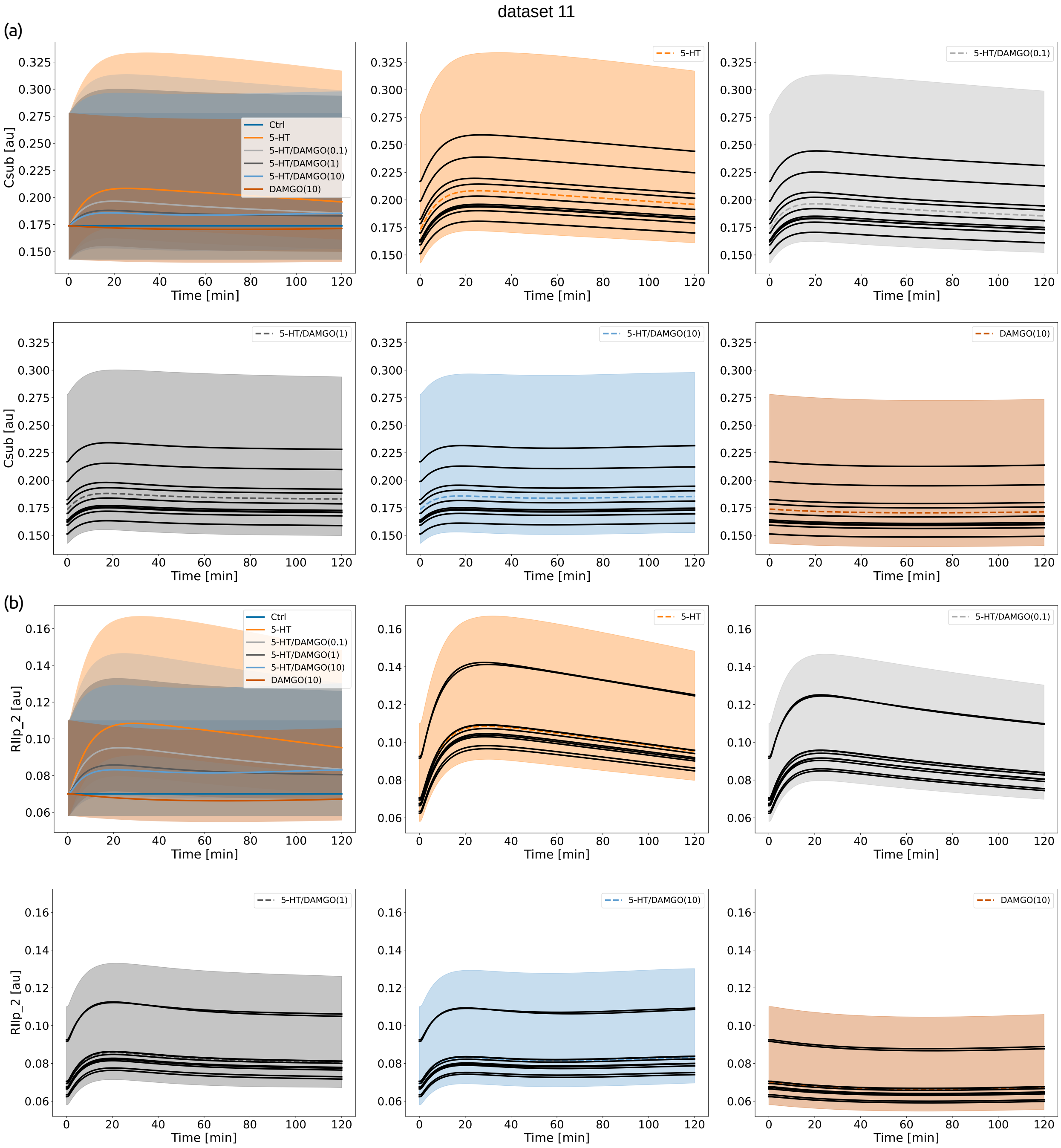


**Fig S4. State variable uncertainty using ensemble-based approach.** Model simulations of selected state variables for LK020_18_LK014_12_MOR_Kinetik_DAMGO_5HT experiment. The state variables (a) C, and (b) pRII we simulated using the ensemble of 2,877 parameters that correspond to statistically indistinguishable objective function values at the 0.01 significance level. The shaded area represents the range of all ensemble simulations.
